# Identifying key biodiversity areas in Mozambique for conservation planning and policy

**DOI:** 10.64898/2026.11.10.764623

**Authors:** Hermenegildo Matimele, Eleutério Duarte, Hugo Costa, Joaquim Campira, Acácio Chechene, Laura Kor, Ara Monadjem, Vanessa Muianga, Armando Sambo, Domingos Sandramo, Celso Sardinha, David Seaman, Jorge J. Sitoe, Joelma Souane, Jonathan Timberlake, Gerson Tomo, Iain Darbyshire, Robert J Smith

## Abstract

The world is experiencing unprecedented rates of biodiversity loss. To reverse this trend, signatories to the Convention on Biological Diversity (CBD) have committed to conserving at least 30% of marine and terrestrial areas by 2030, with an emphasis on sites critical for biodiversity. One mechanism to support this goal is the identification of Key Biodiversity Areas (KBAs) to inform spatial planning and the development of conservation area networks. However, few case studies demonstrate how the KBA process can be implemented in high-biodiversity, low-income countries. We present a national-scale example from Mozambique, where KBAs were delineated across seven taxonomic groups, and outline a post hoc theory of change to support broader national implementation and progress towards CBD targets. We describe how the project established institutional arrangements and training to build technical capacity, highlighting the central role of the Ministry of Land and Environment. We then present the KBA identification process and results, including species Red List assessments that contributed to the identification of four marine and 25 terrestrial KBAs. With results integrated into national policy and legislation, the Mozambique KBA system demonstrates that KBA identification is feasible in lower-income countries and provides a model for broader application.

## 1. Introduction

The world is facing unprecedented rates of biodiversity loss (Butchart et al. 2010; Díaz et al. 2020). In response, the UN Convention on Biological Diversity (CBD) adopted the Kunming-Montreal Global Biodiversity Framework (GBF), which defines 23 targets to “take urgent action to halt and reverse biodiversity loss” (CBD, 2022). Signed by nearly 200 nations, this includes Target 3 to conserve at least 30% of marine and terrestrial areas by 2030.

Commonly referred to as the GBF’s headline 30×30 target, it emphasises the need to focus on areas of particular importance for biodiversity and ecosystem functions and services (CBD, 2022). This is crucial because existing protected area networks are often biased towards remote, economically unproductive areas and charismatic species, leaving many species and ecosystems under-represented (Joppa & Pfaff 2009; Butchart et al. 2015; Venter et al. 2018). Simply expanding conservation area extent to meet percentage targets will therefore not effectively address global species extinction rates or the loss of ecosystems, an issue which has been demonstrated in Mozombique (Richards et al. 2023; Visconti et al. 2019). With research suggesting that more representative protected area networks could increase the number of species under protection by up to 37 times (Venter et al. 2014; Wauchope et al. 2022), a harmonised approach to identifying sites of global biodiversity importance is important(Schaaf & Clamote Rodrigues 2016).

The Key Biodiversity Area (KBA) approach was developed to identify sites that contribute significantly to the global persistence of biodiversity (IUCN 2016). Developed through engagement with taxonomic experts, conservation practitioners, and other stakeholders, the IUCN’s *Global Standard for the Identification of Key Biodiversity Areas* (hereafter KBA Standard) consolidated existing approaches to enable the identification of globally important sites for biodiversity at national or regional levels. It helps overcome known biases in spatial conservation prioritisation (Nogueira et al. 2021) by applying globally consistent criteria with quantitative thresholds for objective, repeatable and transparent analysis (IUCN 2016). This process involves gathering global biodiversity data and combining them with local data and expertise to identify sites that meet threshold proportions of the global distribution of important species and ecosystems.

The GBF monitoring framework includes KBAs as complementary indicators for Targets 1, 2 and 3 (CBD 2024). They therefore play an important role in tracking and supporting progress towards global biodiversity goals (Perino et al. 2022), whose achievement depends on each of the 195 signatory countries fulfilling their commitments under the CBD. KBA identification is most effective when led at the national level, enabling integration into policy, planning, and cross-sectoral decision-making, while fostering national ownership (Perino et al. 2022). However, many high-biodiversity, lower-income tropical countries face constraints in data availability, technical capacity, and financial resources for biodiversity assessment and planning (Hochkirch et al. 2021; Stephenson et al. 2017).

The KBA approach was designed to be applicable under such resource constraints (Meyer et al. 2015). Each potential site is assessed independently against global data, with no requirement for information about other sites in the country (Brooks et al. 2015), thereby allowing for geographically uneven occurrence datasets. In addition, only one species or ecosystem needs to meet any one of the sub-criteria to trigger KBA status, so the process does not depend on surveying for a wide range of taxa. The process involves stakeholders and local, regional and international experts working together to address knowledge gaps (KBA Standards and Appeals Committee 2020). Despite this flexibility, there remains a lack of empirical case studies demonstrating how KBA identification can be implemented at national scale in lower-income countries, particularly with respect to institutional arrangements, capacity development, and integration into policy and practice. This gap is highlighted by an IUCN assessment of protected areas in Africa, which reported in 2020 that only five countries had completed, or were in the process of completing, KBA identification (IUCN ESARO 2020).

To address this gap, we present a national-scale case study from Mozambique. Mozambique is a country with high levels of species richness and endemism (Mittermeier et al. 2004; Darbyshire et al. 2019) and diverse terrestrial, marine and coastal ecosystems (MITADER 2019). It is also one of the poorest countries in the world, with Gross Domestic Product (GDP) per capita among the lowest globally (IMF 2025). To meet its international biodiversity commitments, the Government of Mozambique, through the Ministry of Land and Environment, and with support from conservation partners and donors, led a national-scale process to identify KBAs across multiple taxonomic groups. This required not only the compilation and analysis of biodiversity data, but also the development of institutional structures and technical capacity to support implementation.

In this article, we detail the full KBA process which aimed to: (i) identify sites of global and national importance for biodiversity in Mozambique, and (ii) present a transferable framework for KBA identification and integration into national policy and legislation in comparable high-biodiversity, lower-income countries. We describe the methods used to delineate marine and terrestrial KBAs, document the resulting national network and the biodiversity it supports, and develop a theory of change based on the institutional and capacity-building processes underpinning identification and implementation.

## 2. Methods

### 1.1. Background

Mozambique is located in southeastern Africa, covering a land area of 786,380 km^2^ with a coastline of 2,770 km (MITADER 2015) and approximately 562,000 km^2^ of Exclusive Economic Zone (EEZ). As a signatory to the CBD, Mozambique is committed to contributing to the achievement of GBF targets, including integrating biodiversity and environmental considerations into conservation and development decision-making. This requires robust baseline data on the distribution and conservation status of species, alongside identification of sites of global biodiversity importance. To support this, the Ministry of Land and Environment, in partnership with the Wildlife Conservation Society (WCS), implemented a two-year, USD 250,000 national project from February 2019: *“Red List of Threatened Species and ecosystems, and identification and mapping of Key Biodiversity Areas (KBAs) in Mozambique”* (WCS et al. 2021a).

### 1.2. KBA identification

To assess whether a site can be considered a KBA, the KBA Standard defines 11 globally-consistent criteria in 5 categories, relevant at either the ecosystem or species level (IUCN, 2016). Species-based assessments involve evaluating species data against the KBA criteria to verify whether the site contains a large enough proportion of the trigger species to meet quantitative thresholds (Table 1). While meeting any single criterion or sub-criterion is sufficient for a particular site to trigger KBA status, multiple criteria should be applied across taxonomic groups to strengthen the designation where data allows.

**Table 1:** Key Biodiversity Area criteria and thresholds used in the Mozambique assessment.

| A. Threatened biodiversity |  |
|---|---|
| A1a | ≥0.5% of global population size and ≥5 reproductive units (RU) of a CR/EN species |
| A1b | ≥1.0% of global population size and ≥10 RU of a VU species |
| A1c | ≥0.1% of global population size and ≥5 RU of a species listed as CR/EN due only to past/current decline [= Red List A1, A2, A4 only] |
| A1d | ≥0.2% of global population size and ≥10 RU of a species listed as VU due only to past/current decline [= Red List A1, A2, A4 only] |
| A1e | Effectively the entire population size of a CR/EN species |
| B. Geographically restricted biodiversity |  |
| B1. Individual geographically restricted species | ≥10% of global population size and ≥10 RU of any species |
| B2. Co-occurring geographically restricted species | ≥1% of global population size of each of a number of restricted range species in a taxonomic group: ≥2 species or 0.02% of the total number of species in the taxonomic group, whichever is larger |
| D. Biological processes |  |
| D1. | Demographic aggregations |
| D1b | Site is among largest 10 aggregations of the species |

We applied species-level criteria under categories (A) threatened biodiversity, (B) geographically restricted biodiversity, and (D) biological processes (Table 1), as a national-level ecosystem map was not available at the time. The project approach was based on a successful initiative in Uganda (Plumptre et al. 2019) and adapted to account for local conditions (e.g. Ivanova et al. 2016).

The process followed the structured approach defined in the KBA Standard to ensure transparency and consistency. This included the use of the best available data, expert and stakeholder engagement for validation and context, and documentation and publication of all decisions and thresholds (IUCN, 2016; KBA Standards and Appeals Committee, 2020). The workflow applied to identify KBAs in Mozambique is summarised in Figure 1a and detailed below.

**Figure 1.**
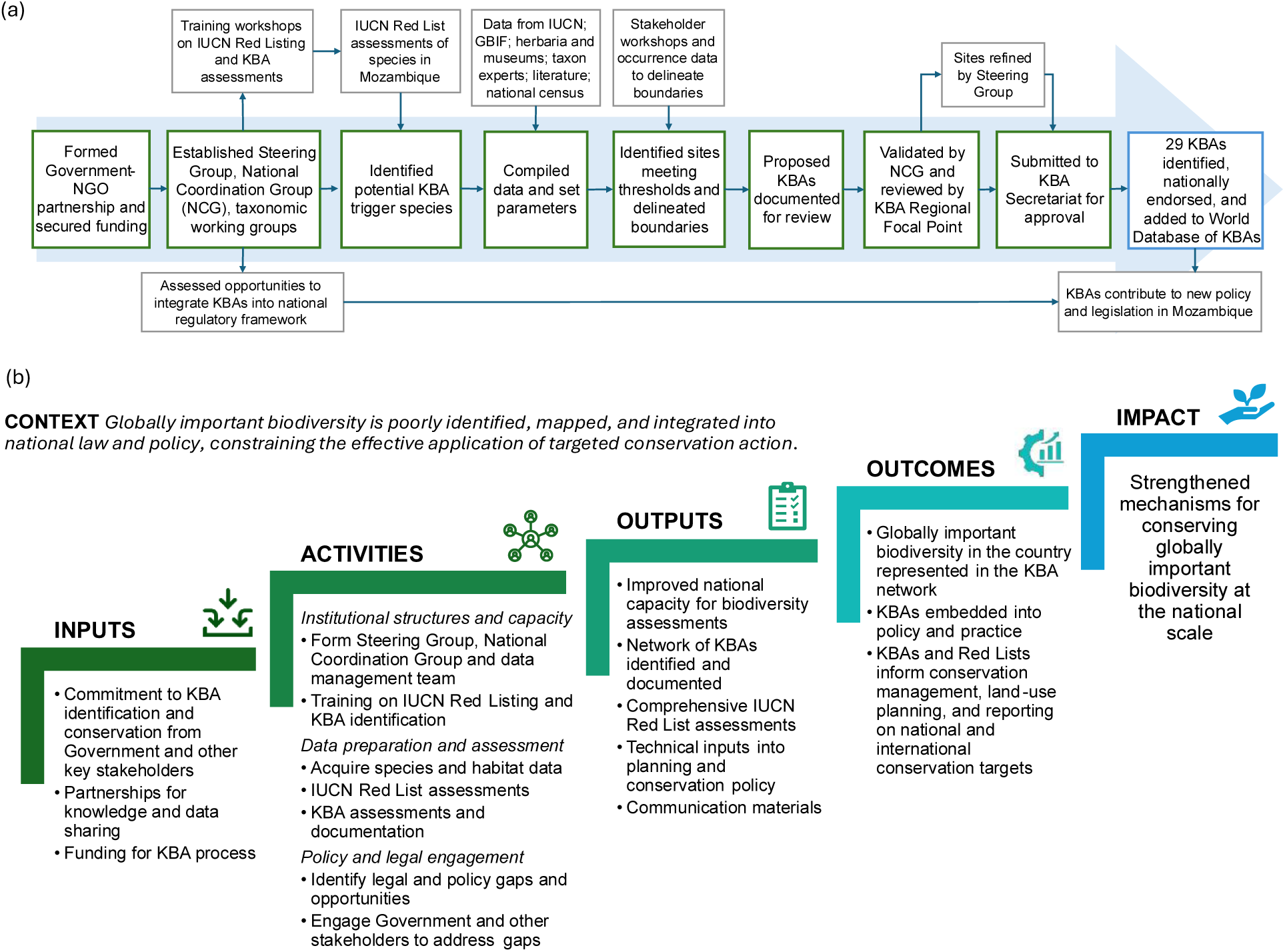
(a) Summary of the workflow used to identify KBAs in Mozambique which represents an operationalisation of the (b) Theory of change for KBA identification in high-biodiversity, lower-income countries based on the Mozambique experience.

### 1.3. Developing enabling conditions

#### 1.3.1. Institutional structures

The Mozambique KBA project began in 2019 with the establishment of institutional arrangements and enabling conditions required to support KBA identification (Figure 1). At the institutional level, a Steering Group was formed to coordinate the process, comprising representatives from WCS Mozambique (including the lead author of this article) and the Ministry of Land and Environment. Technical guidance was provided by the KBA Secretariat, the South African National Biodiversity Institute (SANBI), and the WCS global network.

To provide taxon-specific expertise, the Steering Group established eight working groups focused on amphibians, birds, freshwater fish, invertebrates, mammals, marine biodiversity, plants, and reptiles. Each was led by one or two experts, supported by contributing specialists and a data management assistant responsible for data gathering and preparation.

The Steering Group initiated the process to establish a KBA National Coordination Group. As recommended in IUCN guidance, the role of the National Coordination Group was to facilitate the KBA identification process, help build consensus among stakeholders, oversee national red list assessment processes, validate results at the national level, and support integration into national policy, planning, and decision-making (KBA Standards and Appeals Committee 2020). The National Coordination Group included representatives from 20 national institutions across government, academia, civil society, and the private sector. To ensure government ownership, effective integration of KBAs into national policy, and long-term sustainability, the National Directorate for Environment served as the permanent chair, with the Mozambican Oceanographic Institute as vice-chair. To support the effective operationalisation of the National Coordination Group, a management committee was established and secretariat formed, with designated focal points for KBA review, Red List assessments, and data management.

#### 1.3.2. Capacity building and species red listing

Capacity building was integrated throughout the process to support ensure accurate application of the KBA Standard and long-term national ownership (Figure 1a). The Steering Group organized a national training workshop for academics, scientists, taxon experts, and stakeholders to introduce KBA criteria, thresholds, and their application to Mozambique-specific datasets.

To enable the application of Criterion A1 (threatened biodiversity), Red List data were required. Red List assessments for endemic and near-endemic plants already existed, primarily as part of the Tropical Important Plant Areas programme (Darbyshire et al. 2023) and through the IUCN-SSC Southern African Plant Specialist Group (IUCN, no date). However, gaps remained for a number of faunal taxonomic groups. The National Coordination Group identified this as a critical limitation and prioritised Red List assessments for endemic and near-endemic taxa lacking global assessments.

The Steering Group held a regional training workshop in 2019 for 41 taxonomic experts from Mozambique, Eswatini, and South Africa. This led to Red List assessments for 67 species across amphibians, freshwater fish, invertebrates, and reptiles, based on collated occurrence and ecological data (Duarte & Matimele, 2020).

### 1.4. KBA trigger species

We compiled a list of species likely to meet the criteria and thresholds of the KBA Standard and therefore trigger KBA status at a given site (hereafter “trigger species”) (IUCN 2016). The criteria and sub-criteria applied are listed in Table 1.

Species likely to trigger Sub-Criterion A1 (threatened biodiversity) were identified from those that had IUCN Red List assessments and with data available for download from the Red List web site (Table S-1). These datasets included species locality records, range maps, and estimates of the number of mature individuals (IUCN 2016).

For Sub-Criterion B1 (geographically restricted biodiversity), we focused on nationally endemic species whose global range is wholly restricted to Mozambique (Kraus et al. 2023). Data for endemic and near-endemic species were primarily obtained from taxon experts, supplemented by data from herbaria, museum databases, and literature. Global datasets such as the Global Biodiversity Information Facility were screened by taxonomic experts and ultimately excluded, as records were inconsistent or did not improve spatial resolution compared to the curated national and local datasets. Sub-Criterion B2 focuses on range restricted species, defined as having a global distribution of ≤10,000 km^2^ (IUCN 2016). Most nationally endemic species qualified under this criterion, and we also included relevant near-endemic species, defined as species whose range distribution is not entirely within Mozambique but is ≤10,000 km^2^ (Darbyshire et al. 2019).

Under Criterion D (Biological Processes), species exhibiting demographic aggregations (Sub-Criterion D1) were assessed with a focus on bird and mammal species, as they are taxonomic groups with broader ranges and generally better-documented life history patterns. We obtained global datasets from the IUCN Red List, while national occurrence and biology of these species in Mozambique were obtained from taxon experts and national population census reports from the National Administration of Conservation Areas. For these species, assessments focused on identifying management units containing sufficient proportions of global populations to meet thresholds to trigger KBA status.

### 1.5. KBA assessment

#### 1.5.1. Site identification

KBA assessments evaluated whether sites contained sufficient proportions of trigger species to meet quantitative thresholds (Table 1). Where population data were unavailable, alternative parameters were used, including extent of suitable habitat, range size, and number of localities (IUCN, 2016; KBA Standards and Appeals Committee 2020).

Considering this, our dataset allowed the application of Sub-Criteria A1: threatened species, B1: individual geographically restricted species, B2: co-occurring geographically restricted species, and D1: demographic aggregation (IUCN 2016).

With over 60% of our data consisting of georeferenced occurrences, assessments were primarily based on the number of localities, defined as “a discrete population to the extent this can be inferred, given the degree of habitat fragmentation and what is known about the dispersal capabilities of the species” (IUCN 2022). We used the number of locations reported in IUCN Red List assessments as a proxy for localities, as defined in KBA guidelines (IUCN, 2022). This was considered appropriate given the restricted distributions and habitat fragmentation of most species assessed.

#### 1.5.2. Delineation and validation

We delineated geographic boundaries for potential KBA sites using a combination of desktop studies, expert workshops, and consultation with stakeholders. The KBA Standard requires that site boundaries should be both ecologically relevant and suitable for management as one single unit (IUCN, 2016). We derived initial site boundaries by overlaying distributional data for the KBA trigger species at each site to determine suitable ecological boundaries (KBA Standards and Appeals Committee 2020). For species with limited data, distributions were inferred based on expert knowledge of habitat requirements. Where sites overlapped with existing biodiversity areas (e.g. Important Bird and Biodiversity Areas or Alliance for Zero Extinction sites), consultation was undertaken to build consensus with proposers of the existing site and agree on boundary modification where necessary.

To ensure practical boundaries, KBA guidelines encourage consultation to account for existing management units (e.g. protected areas) and consider land ownership and stakeholder views (KBA Standards and Appeals Committee 2020). Coordinated by the Steering Group, we held a workshop involving 61 participants from 33 institutions, consisting of a wide range of taxonomic experts and civil society stakeholders, to discuss and create consensus on the delineation of boundaries for each potential KBA site. Where proposed KBAs overlapped with an existing PA, boundary delineation followed the borders of that PA. Following delineation, sites were reassessed to confirm that thresholds were met.

The Steering Group submitted the KBA draft assessments to the National Coordination Group for review, who then evaluated the identification and delineation process, ensuring incorporation of expert and stakeholder input. Validated and revised proposals were forwarded to the KBA Regional Focal Point for approval before submission to the KBA Secretariat for final validation. Of 30 proposed sites, 29 were approved and included in the World Database of KBAs.

#### 1.5.3. Integrating KBAs into national policy and legal framework

A core aim of the project was to support the integration of KBAs into national policy and legislation, including as avoidance areas for development projects. Government institutions were therefore strongly represented in the National Coordination Group, and targeted awareness-raising and capacity-building activities were implemented throughout the project for a range of stakeholders (Figure 1a).

The project team assessed management and protection options for delimited KBAs and identified entry points for embedding them, and the Red List of Threatened Species, within spatial planning frameworks and environmental legislation. This included their use as avoidance areas for development. A suite of communication and dissemination materials were produced by the Steering Group, including technical reports, infographics, an online GIS, a story map, and a summary brochure documenting the process. Results were presented to a broad audience in a workshop led by the Ministry of Land and Environment, supporting the formal recognition of KBAs as a key conservation planning tool.

#### 1.5.4. Data analysis

Descriptive statistics were used to summarise key parameters, including the number of species triggering KBAs, disaggregated by taxonomic group, level of endemism, and threat. We also analysed the criteria responsible for triggering each KBA and quantified the extent of KBA sites that are protected versus those lacking formal protection. Data management, analyses and visualisation were conducted in R v.4.3.0 (R Core Team 2022) and Excel, with mapping and spatial analysis in QGIS 3.30 QGIS Development Team (2023).

### 1.6. Theory of change

A theory of change involves defining the long-term goals for a project or strategy and mapping out the necessary actions and preconditions for success. We developed a post hoc theory of change to describe the pathways through which national-scale KBA identification can support conservation outcomes in high-biodiversity, lower-income countries. Drawing on the Mozambique process (Figure 1a) and established approaches to theory of change development in conservation (Rice et al., 2020), we defined key enabling conditions, activities, outputs, and outcomes required for effective implementation and integration into policy and practice.

## 3. Results

The Mozambique KBA identification process was implemented over a two-year period and resulted in the identification and mapping of 25 terrestrial and four marine KBAs across the country (Table 2; Figure 3). The process involved three major workshops and numerous consultations with over 100 national, regional and international experts, including representatives from 20 national institutions. It established the KBA National Coordination Group, which oversaw IUCN Red Listing and KBA identification, the results of which are detailed in this section.

**Table 2:** List of Key Biodiversity Areas of Mozambique, including the province in which they are located, the number of trigger species they contain and the KBA Sub-Criteria of these trigger species.

| ID | KBA name | KBA area (km<sup>2</sup>) | Province | Trigger species | KBA Sub-Criteria of trigger species |
|---|---|---|---|---|---|
| 1 | Njesi Plateau | 1,996 | Niassa | 2 | A1a; A1e, B1 |
| 2 | Niassa Special Reserve | 42,708 | Niassa | 7 | A1a; A1b; A1c; A1d, B1 |
| 3 | Palma | 4,556 | Cabo Delgado | 17 | A1a, A1b, A1e, B1, B2 |
| 4 | Vamizi | 87 | Cabo Delgado | 1 | D1b |
| 5 | Quiterajo | 3,064 | Cabo Delgado | 15 | A1a, A1b, A1e, B1, B2 |
| 6 | Taratibu | 25 | Cabo Delgado | 2 | A1a; A1e, B1 |
| 7 | Eráti | 148 | Nampula | 1 | A1a, B1 |
| 8 | Matibane Forest Reserve | 109 | Nampula | 1 | A1a, B1 |
| 9 | Ribáuè-Mphalwe | 265 | Nampula | 15 | A1a; A1b; A1e; B1; B2 |
| 10 | Mount Inago | 326 | Nampula | 7 | A1a, A1e, B1, B2 |
| 11 | APAIPS | 2,512 | Zambezia | 1 | A1a, B1 |
| 12 | Mount Namuli | 53 | Zambezia | 30 | A1a; A1b; A1e; B1; B2 |
| 13 | Mount Mabu | 61 | Zambezia | 17 | A1a, A1b, B1; B2 |
| 14 | Mount Chiperone | 36 | Zambezia | 4 | A1b, B1, B2 |
| 15 | Derre Forest Reserve | 3,984 | Zambezia | 2 | A1b, B1 |
| 16 | Tchuma-Tchato Cahora Bassa Lake | 38,180 | Tete | 2 | A1a, A1b, A1c, A1d |
| 17 | Serra Choa | 516 | Manica | 1 | A1b |
| 18 | Machipanda | 756 | Manica | 2 | A1b |
| 19 | Chimanimani National Park | 2,371 | Manica | 41 | A1a; A1b; A1e; B1 |
| 20 | Gorongosa and Marromeu Complex | 23,088 | Sofala | 21 | A1a, A1b, A1c, A1d, A1e, B1 |
| 21 | Inhassoro-Vilankulos | 5,357 | Inhambane | 5 | A1a; A1b; B1 |
| 22 | Great Bazaruto | 5,236 | Inhambane | 7 | A1a, A1b, A1c, A1d, B1 |
| 23 | Tofo | 342 | Inhambane | 4 | D1b |
| 24 | Chongoene | 33 | Gaza | 1 | A1a, B1 |
| 25 | Manhiça-Bilene | 2,070 | Gaza | 3 | A1a, A1b, A1e, B1 |
| 26 | Matutuine | 195 | Maputo | 2 | A1b, B1 |
| 27 | Licuáti Forest Reserve | 141 | Maputo | 6 | A1a, A1b, B1 |
| 28 | Maputo National Park | 1,040 | Maputo | 5 | A1a; A1b; B1 |
| 29 | Ponta do Ouro Partial Marine Reserve | 698 | Maputo | 3 | A1b; A1c, D1b |

### 1.7. KBA trigger species, endemism and IUCN red list status

The Mozambique KBA analysis comprised 180 species (Table S-1), consisting of 102 plants, 26 reptiles, 14 fish (8 freshwater and 6 marine), 11 birds, 11 insects, 8 amphibians, and 8 mammals (6 terrestrial and 2 marine). Of these, 128 are endemic to Mozambique, most of which are plants (83 species) and reptiles (25 species). Endemism levels varied by taxonomic group, ranging from 12.5% for mammals to 96.2% for reptiles (Figure 2a).

**Figure 2:**
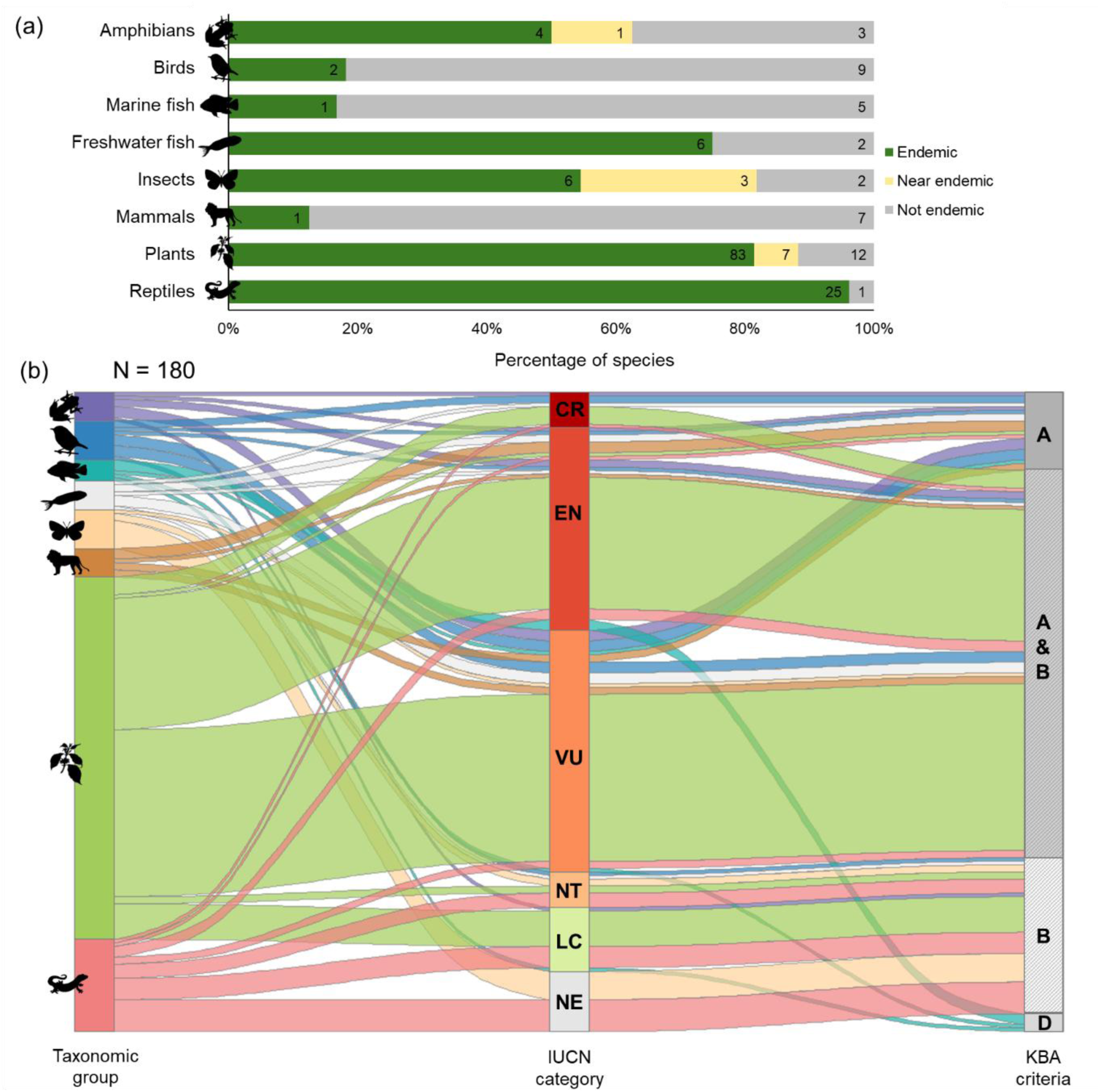
(a) Endemism levels per taxonomic group; (b) species global conservation status per taxonomic group based on the IUCN red list of threatened species and the KBA criteria triggered.

Conservation assessments were available for 163 of the trigger species on the IUCN Red List, leaving 17 species of fish, insect and reptile species not evaluated. Of the assessed species, 75.0% are listed as Critically Endangered, Endangered or Vulnerable, with proportions ranging from 89.2% for plants and 100.0% for the assessed mammals (Figure 2b).

### 1.8. KBA criteria and trigger species

The 180 species triggered three high-level KBA Criteria: (A) Threatened biodiversity, (B) Geographically restricted biodiversity, and (D) Biological processes (Table 1).

Criterion A was triggered by a total of 133 species, with 22 of these species meeting no additional criteria (Figure 2b). Of these, 37.2% triggered Sub-Criterion A1b due to their Vulnerable status, while 34.4% triggered A1a because they are either Critically Endangered or Endangered. Fewer species met the other sub-criteria under Criterion A: 9.4% triggered A1e as Critically Endangered or Endangered with their global population confined to a single site, 3.3% triggered A1c due to their Critically Endangered or Endangered status and ongoing or past population decline, and 2.2% triggered A1d due to their Vulnerable status and ongoing or past population decline.

Criterion B was triggered by 153 species, with 42 of them triggering no other criteria (Figure 2b). Of these, 84.4% met Sub-Criterion B1 because they are geographically restricted to one site across their life stages, and 15.6% met B2 as they were found in sites with multiple co-occurring geographically restricted species (Table 1).

Only five species triggered Criterion D, all of them not triggering any other criteria. They all met Sub-Criterion D1b, found at sites where the population of mature individuals is representative of their 10 largest aggregations worldwide (Table 1).

### 1.9. Sites of global importance and overlap with protected areas

The 29 KBAs identified in Mozambique cover a total of 139,947 km^2^. This consisted of 25 terrestrial sites, which cover 17% of the country’s continental and island territory, and four marine sites, which cover 1% of the EEZ (Figure 3). Of these, 24 sites contained species triggering both Criteria A and B, one site triggered Criteria A and D, two sites triggered Criterion A only, and two sites triggered Criterion D only (Table 2; Figure 3).

**Figure 3:**
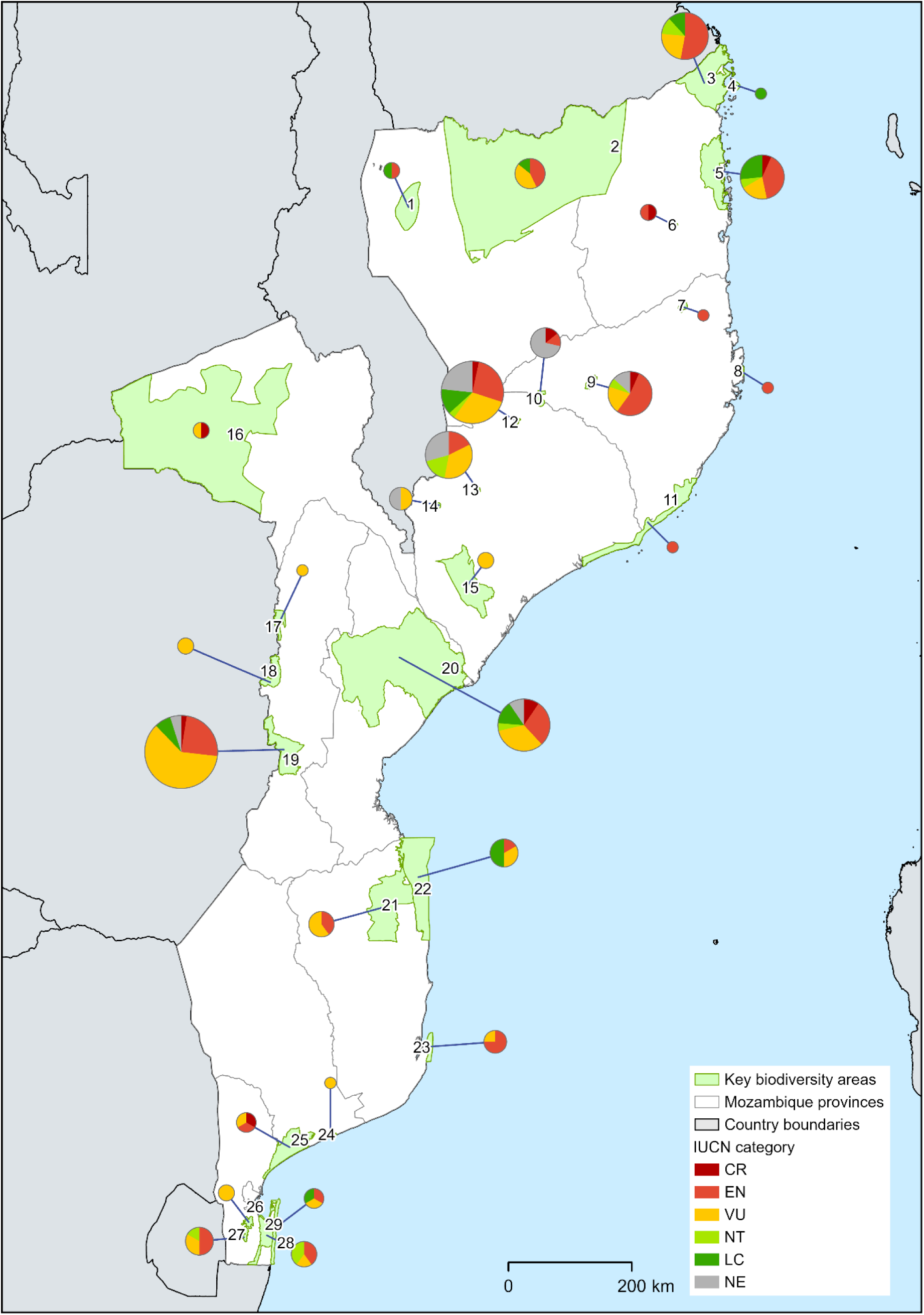
The 29 Key Biodiversity Areas identified in Mozambique. Numbers refer to the site IDs listed in Table 2, while pie charts are proportionate to the number of KBA trigger species in each site and their IUCN threat status.

Criterion A1e was triggered in 10 sites, each containing the entire global population of at least one threatened species (Critically Endangered or Endangered). Across the network, 72% of sites included species triggering Sub-Criterion A1a, while 10% included species triggering A1d (Vulnerable with ongoing or past declines). Criterion B was triggered in 23 sites through Sub-Criterion B1 (geographically restricted species), and in three sites via B2 (co-occurring restricted species). Criterion D1b was triggered in five species across three sites, representing some of the largest known global aggregations.

Overall, 85% of KBA area overlaps partially or fully with existing designated conservation areas. This includes overlaps with protected areas (48% of KBA sites), forest reserves (20%), and internationally designated sites such as Ramsar sites, World Heritage Sites, and Biosphere Reserves (17%).

### 1.10. Mainstreaming of KBAs into national policy

Following approval by the KBA Secretariat, the KBAs were formally endorsed by the Minister of Land and Environment and publicly launched at a national event attended by 57 in person and over 100 remote participants from government, private sector, academia, and civil society. Subsequent engagement through the Steering Group and National Coordination Group facilitated integration of KBAs into six national policy and legislative instruments.

These include spatial planning frameworks such as the National Territorial Development Plan (Resolution No. 7/2021) and the Marine Spatial Plan (Resolution No. 63/2024), both of which identify KBAs as areas where development impacts should be avoided.

Sectoral legislation similarly incorporates KBAs, including the National Forest Law (Law No. 17/2023) and associated regulations (Decree No. 78/2024), which recognise forests within KBAs as conservation priority areas. The Decree for the Protection of Birds and their Habitats (Decree No. 51/2021) and the Biodiversity Offsets Directive (Ministerial Order No. 55/2022) further embed KBAs within regulatory frameworks by defining them as avoidance areas and preferred offset receiving sites.

### 1.11. Theory of change

A post hoc theory of change was derived from the Mozambique KBA process (Figure 1b), linking national context, project inputs, activities, outputs, and outcomes to inform the process in other high-biodiversity, lower-income countries. Inputs included government commitment, multi-stakeholder partnerships, and institutional and financial support. Activities comprised the establishment of governance structures (Steering Group and National Coordination Group), capacity building in Red Listing and KBA criteria, targeted Red List assessments, KBA identification and documentation, and sustained policy engagement.

These activities generated outputs described in Sections 3.1–3.4, including a national KBA network and expanded Red List coverage. Outcomes included improved representation of globally important biodiversity within national planning frameworks and formal integration of KBAs into key policy and legislative instruments (Section 3.5).

## 4. Discussion

### 1.12. Mozambique’s KBAs

Mozambique is characterized by high species richness and endemism, overlapping with three regional biodiversity hotspots (Marchese 2015) and six centres of plant endemism (Darbyshire et al. 2019). As with many low-income tropical countries, significant data gaps exist across taxonomic groups (Meyer et al. 2015; Monadjem et al. 2010; Neves et al. 2019; Stephenson & Stengel 2020; Tolley et al. 2016). In this context, and in the absence of resources to generate new primary biodiversity datasets, this study built on recent national and regional initiatives focused on consolidating and curating existing biodiversity information (e.g. Malatesta et al. 2023).

The identification of a network of KBAs in Mozambique based on multiple taxonomic groups shows that the KBA Standard can be applied effectively even in data-limited settings. Viewed in comparison to countries with more complete biodiversity datasets and established national KBA processes (e.g. SANBI, 2024), our results suggest that successful application depends on coordination, institutional capacity, and broad collaboration rather than data completeness alone (Langhammer et al., 2007; IUCN, 2016). The framing of these enabling conditions within a theory of change (Figure 1b) provides a structured model for identifying KBAs in similar contexts. This is important to support the GBF’s 30×30 target which goes beyond coverage, emphasising the need for area-based conservation to incorporate areas of particular importance for biodiversity (Visconti et al. 2019; CBD 2022).

### 1.13. Implications of data availability

The KBA Standard was designed for use in a range of contexts, including data poor conditions. It allows site-by-site assessment using global species data and the application of any one or more of the criteria (Brooks et al. 2015; KBA Standards and Appeals Committee 2020). This flexibility proved important in this study by allowing different criteria to contribute in complementary ways. Some species triggered KBAs under Criterion A (threatened biodiversity), others under criterion B (restricted-range species), and a small number under Criterion D (biological aggregations). Criterion D, although applied to fewer species, was critical for identifying ecologically important aggregations that would otherwise not be captured under threat- or range-based approaches alone. This reinforces the value of a multi-criteria framework for capturing complementary aspects of biodiversity importance (IUCN 2016; Perino et al. 2022).

Despite this flexibility, the results also reflect how data availability and sampling bias influence KBA identification (Handley et al. 2023). Most KBAs were triggered by relatively few species, with several sites containing only one or two trigger species (Table 2). While this may reflect genuine patterns of endemism and rarity, it also likely reflects uneven sampling effort and data availability. Under-detection of species is a known challenge in biodiversity datasets and can result in false absences and biased estimates of range distribution and population size (Chen et al. 2013; Lomba et al., 2010; Moore et al. 2011; Zhang et al. 2014). This is particularly relevant for rare and threatened species (Alexander et al. 2012). In Mozambique, sampling was generally highest in existing protected areas, which may have contributed to the strong overlap between KBAs and these protected sites.

Sparsity of records may also reflect true biological rarity (Rivers et al. 2011; Roberts et al. 2016) and it is possible that the sites contain other important species for which data are not available. This is particularly pertinent given that an additional 15 sites were identified as likely potential KBAs if more data were available (WCS, 2021a), and some of these sites overlap with Important Plant Areas that have been identified since the KBA project ended (Darbyshire et al. 2023; Richards et al. 2023). Marine ecosystems remain particularly under-represented, reflecting both data scarcity and historical research bias, with existing literature highlighting the need for more in-depth assessments in marine hotspot areas (Pereira et al. 2014).

Taken together, more comprehensive biodiversity data will likely reveal additional KBAs in Mozambique. Alongside the KBA Standard’s guidance to re-evaluate sites every 8–12 years to incorporate new data and reduce uncertainty (IUCN, 2016), we therefore recommend increasing efforts to gather such data across the country and conduct a comprehensive national biodiversity assessment, both in terrestrial and marine realms.

Given data limitations, this study demonstrated the importance of interpretation, expert input, and institutional coordination for the application of the KBA Standard at a national scale.

Assessing against Criteria such as B1 and D1 often required expert-derived estimates of population size, locality structure, or aggregation behaviour, highlighting the importance of expert knowledge in compensating for data gaps (KBA Standards and Appeals Committee 2020). The structured and transparent assessment process helps ensure consistency and repeatability across taxa and sites when using stakeholder inputs.

### 1.14. Governance and integration into law and policy

Integration of biodiversity priorities into national legal and policy frameworks is an important step in translating global commitments under the CBD into nationally implementable actions (Perino et al. 2022; CBD 2024). KBA status does not prescribe governance or management arrangements (IUCN, 2016). However, the process of site delineation considers current and potential management, so that when trigger species or ecosystems fall within recognised conservation areas, the boundary of that area is often used to delineate the KBA. In Mozambique, this resulted in substantial overlap between KBAs and protected areas, with 12 fully protected and 7 partially protected sites. Such overlap facilitates immediate integration into existing governance systems, although management plans may require updating to explicitly address KBA trigger features and the potential benefits of this are based on the assumption that existing protected areas are effective.

For KBAs outside the protected area network, management must reflect the social and political context of the area they are located in (Wauchope et al. 2022). Options to bring KBAs into formal conservation management in Mozambique include establishing new PAs that are co-managed with, or supported by, conservation NGOs or philanthropic organisations through technical and financial assistance, with both models already successfully applied in the country. Examples include the joint management of Gorongosa National Park through a long-term public-private agreement between the Government of Mozambique and the Carr Foundation, an NGO founded by a US entrepreneur and philanthropist (Gorongosa National Park, 2026).

Some KBAs may also be suitable for recognition as other effective area-based conservation measures (OECMs) – sites that are not PAs but are managed in ways that have positive biodiversity conservation outcomes (IUCN-WCPA Task Force on OECMs 2019). In Mozambique, existing designations which could be considered OECMs include forest reserves, hunting areas, sacred forests, and community-based ecotourism initiatives, should sites be found to meet the OECM criteria. While identification as OECMs can broaden stakeholder engagement, improve recognition of existing community conservation efforts, and contribute to the achievement of the GBF’s 30×30 target, their ultimate effectiveness depends on governance quality and sustained incentives rather than designation alone (Jonas et al. 2024).

Financing mechanisms, including biodiversity offsets and donor funding, offer potential opportunities to support KBA management. Several international donors now prioritise KBAs as a consideration in their funding decisions (KBA Partnership 2025). Following the KBA process in Mozambique (Figure 1a), the national offsets directive has integrated KBAs by identifying them as avoidance areas and preferred offset receiving sites, creating opportunities to channel offset finance into KBA enhancement and long-term protection. This aligns with the mitigation hierarchy and “no net loss” principles, which require credible baselines, avoidance of impacts, measurable biodiversity gains, and durable legal security for offset sites (Maron et al., 2018). This integration of KBAs into Mozambican law and policy reflects the importance of institutional coordination and systemic stakeholder engagement, as captured in the project’s theory of change (Figure 1b).

### 1.15. KBAs within a systematic conservation planning framework

While KBA identification provides a robust standardised approach for recognising sites of global biodiversity importance, it does not explicitly account for national priorities (Liang et al. 2016; Mori et al. 2017; Brockerhoff et al. 2017; Dobson et al. 2022), nor does it incorporate requirements for ecological connectivity needed to maintain landscape functionality and facilitate species’ responses to climate change (Thomas and Gillingham, 2015). It is therefore important to recognise that additional considerations should be made when designing effective conservation area networks (Smith et al. 2019; Handley et al. 2023). Of particular relevance are the CARE principles of systematic conservation planning, comprehensiveness, adequacy, representativeness, and efficiency, which aim to ensure that conservation area networks function as coherent and effective systems (Margules & Pressey 2000). This is important in the context of the GBF 30×30 target, which not only highlights the importance of the coverage of conservation area networks, but also their ecological representativeness and connectivity (CBD 2022; Visconti et al. 2019).

In Mozambique, incorporating KBAs into systematic conservation planning could support the development of more effective conservation networks (Margules & Pressey 2000; Smith et al. 2019). Various ecosystems remain relatively understudied in the country, potentially explaining why few grasslands, woodlands, or savanna systems were identified as KBAs. In addition, several sites support nationally important populations of trigger species but did not meet global KBA thresholds. Applying an SCP approach would ensure that KBAs inform decisions on where to prioritise conservation action, while helping to ensure ecological representation, delivering multifunctional landscapes that maintain connectivity (Kaim et al. 2017; Iwamura et al. 2018; Cunningham et al. 2021) and account for the local context and availability of funding from national governments, donors and agencies (Joseph et al. 2009; Bolam et al. 2019). Such a pathway, from credible site identification to network design, financing, and implementation, reflects the outputs to outcomes pathway articulated in our theory of change (Figure 1b).

### 1.16. Synthesis and implications

The KBA methodology was designed as an umbrella approach encompassing all taxonomic groups and ecosystems to support national environmental decision making and foster government ownership (Perino et al. 2022). In Mozambique, it has enabled the development of a standardised, nationally coordinated process for identifying biodiversity priorities and informing governance and planning, strengthening the country’s ability to reconcile economic development with biodiversity conservation. This has already produced positive results, with two KBAs incorporated into the new Palma-Quirimbas Environmental Protection Area in 2024 (Conselho de Ministros, 2024). This is particularly significant in supporting less charismatic species which may otherwise be overlooked, allowing for their targeted conservation and monitoring within KBA sites (KBA Standards and Appeals Committee 2020). As a signatory to the CBD, the country is now better equipped to update its National Biodiversity Strategy and Action Plan and reporting on progress internationally, particularly for targets related to KBAs and the Red List of species and ecosystems.

This case study from Mozambique represents one of the first national-scale, multi-taxon applications of the KBA Standard in a lower-income country and demonstrates that rigorous, globally consistent site identification is feasible under substantial data constraints. Key enabling factors included strong institutional coordination, capacity building, and sustained stakeholder engagement through a National Coordination Group. The process demonstrates how KBAs can move beyond technical designation to become operational in conservation policy and practice – linking biodiversity evidence directly to spatial planning, legal frameworks, and investment decisions. These findings provide a transferable model and theory of change for other countries implementing the KBA Standard, particularly those seeking to meet the targets of the Global Biodiversity Framework through nationally led, policy-relevant biodiversity planning.

## Author contributions

Hermenegildo Matimele: conceptualisation; data curation; formal analysis investigation; methodology; validation; writing – original draft preparation. Hugo Costa: conceptualisation, funding acquisition, methodology, project administration, supervision, writing – review & editing. Eleutério Duarte and Acácio Chechene: conceptualisation; data curation; formal analysis; investigation; methodology; validation. Armando Sambo, Celso Sardinha, Domingos Sandramo, Gerson Tomo, Joelma Souane, Jorge J. Sitoe, Joaquim Campira and Vanessa Muianga: data curation; formal analysis. Laura Kor: visualisation; writing – original draft preparation; writing – review and editing. David Seaman: writing – review and editing. Robert Smith, David Roberts, Iain Darbyshire: conceptualisation; supervision; writing – review and editing.

## Acknowledgements

The authors would like to thank the United States Agency for International Development (USAID), through the SPEED+ Project *“Supporting the Policy Environment for Economic Development”*, alongside the donors: Agence Française de Développement (AFD), Fonds Français pour l’Environnement Mondial (FFEM), Fondation MAVA, NORAD, and The Tiffany & Co. Foundation. We extend thanks to the SECOSUD-II Project for providing data management assistants and all institutions and individuals who contributed data, logistical support, and expertise. LK, DS and RJS were supported by Research England’s Expanding Excellence in England (E3) Fund, UK Research and Innovation.

## Data availability statement

The list of species used as the basis for KBA identification is openly available from Figshare: 10.6084/m9.figshare.32882027.

## References

Alexander HM, Reed AW, Kettle WD, Slade NA, Bodbyl Roels SA, Collins CD, Salisbury V. 2012. Detection and Plant Monitoring Programs: Lessons from an Intensive Survey of Asclepias meadii with Five Observers. PLoS ONE 7:e52762.

Ament JM, Cumming GS. 2016. Scale dependency in effectiveness, isolation, and social-ecological spillover of protected areas. Conservation Biology 30:846–855. [Wiley, Society for Conservation Biology].

Bolam FC, Grainger MJ, Mengersen KL, Stewart GB, Sutherland WJ, Runge MC, McGowan PJK. 2019. Using the Value of Information to improve conservation decision making. Biological Reviews 94:629–647.

Brockerhoff EG et al. 2017. Forest biodiversity, ecosystem functioning and the provision of ecosystem services. Biodiversity and Conservation 26:3005–3035.

Brooks TM, Cuttelod A, Faith DP, Garcia-Moreno J, Langhammer P, Pérez-Espona S. 2015. Why and how might genetic and phylogenetic diversity be reflected in the identification of key biodiversity areas? Philosophical Transactions of the Royal Society B: Biological Sciences 370:20140019. Royal Society.

Butchart SHM et al. 2010. Global Biodiversity: Indicators of Recent Declines. Science 328:1164–1168. American Association for the Advancement of Science.

Butchart SHM et al. 2015. Shortfalls and Solutions for Meeting National and Global Conservation Area Targets. Conservation Letters 8:329–337.

CBD. 2022. First draft of the post-2020 global biodiversity framework:12.

CBD 2024 Monitoring framework for the Kunming-Montreal Global Biodiversity Framework. Subsidiary Body on Scientific, Technical and Technological Advice. Twenty-sixth meeting, Nairobi, 30 March 2024. Available at : https://www.cbd.int/doc/c/8fad/193d/e30d0be38e537019cdbefb35/sbstta-26-02-en.pdf

Chen G, Kéry M, Plattner M, Ma K, Gardner B. 2013. Imperfect detection is the rule rather than the exception in plant distribution studies. Journal of Ecology 101:183–191.

Conselho de Ministros, 2024. Boletim da República Decreto No. 71/2024. I Série Número 174. Imprensa Nacional ee Moçambique, E.P. Available at: https://archive.gazettes.africa/archive/mz/2024/mz-government-gazette-series-i-dated-2024-09-05-no-174.pdf

Cunningham CA, Crick HQP, Morecroft MD, Thomas CD, Beale CM. 2021. Translating area-based conservation pledges into efficient biodiversity protection outcomes. Communications Biology 4:1–5. Nature Publishing Group.

Darbyshire I et al. 2019. The endemic plants of Mozambique: diversity and conservation status. PhytoKeys 136:45–96.

Darbyshire I et al. 2023. The important plant areas of Mozambique. Royal Botanic Gardens, Kew.

Di Marco M, Butchart SHM, Visconti P, Buchanan GM, Ficetola GF, Rondinini C. 2016. Synergies and trade-offs in achieving global biodiversity targets. Conservation Biology 30:189–195.

Díaz S et al. 2020. Set ambitious goals for biodiversity and sustainability. Science 370:411–413. American Association for the Advancement of Science.

Dobson A et al. 2022. Savannas are vital but overlooked carbon sinks. Science 375:392–392. American Association for the Advancement of Science.

Duarte E and Matimele H (2020). Red List of Threatened Species, Identification and Mapping of Key Biodiversity Areas (KBAs) in Mozambique. Report with Initial Red List of Threatened Species for Amphibians, Reptiles, Freshwater Fish and Lepidoptera.

MITADER, June 2020. Available from: https://sibmoz.gov.mz/key-biodiversity-areas/

Gorongosa National Park, 2026. Our Story. Available from: https://gorongosa.org/ (accessed July 2, 2026)

Handley JM, Harte E, Stanworth A, Poncet S, Catry P, Cleminson S, Crofts S, Dias M. 2023. Progressing delineations of key biodiversity areas for seabirds, and their application to management of coastal seas. Diversity and Distributions 29:123–142.

Hochkirch, A., Samways, M.J., Gerlach, J., Böhm, M., Williams, P., Cardoso, P., Cumberlidge, N., Stephenson, P.J., Seddon, M.B., Clausnitzer, V., Borges, P.A.V., Mueller, G.M., Pearce-Kelly, P., Raimondo, D.C., Danielczak, A. and Dijkstra, K.-D.B. 2021. A strategy for the next decade to address data deficiency in neglected biodiversity. Conservation Biology, 35: 502–509. 10.1111/cobi.13589

IMF. 2025. World Economic Outlook Database, October 2025 Edition. International Monetary Fund. Available from: www.imf.org/external/datamapper/PPPPC@WEO/OEMDC/ADVEC/WEOWORLD (accessed January 13, 2026)

IUCN. No date. IUCN SSC Southern African Plant Specialist Group. Available from: https://iucn.org/our-union/commissions/group/iucn-ssc-southern-african-plant-specialist-group (accessed January 13, 2026)

IUCN. 2012. IUCN Red List categories and criteria, version 3.1, second edition. IUCN, Gland, Switzerland and Cambridge, UK. Available from https://portals.iucn.org/library/node/10315 (accessed February 12, 2023).

IUCN. 2016. A global standard for the identification of key biodiversity areas. Available from https://portals.iucn.org/library/sites/library/files/documents/2016-048.pdf (accessed November 9, 2022).

IUCN. 2022. The IUCN Red List of Threatened Species. Version 2022-1. Available from https://www.iucnredlist.org/en (accessed November 10, 2022).

IUCN ESARO. 2020. The state of protected and conserved areas in Eastern and Southern Africa. State of Protected and Conserved Areas Report Series No. 1. Nairobi, Kenya: IUCN ESARO.

IUCN-WCPA Task Force on OECMs. 2019. Recognising and reporting other effective area-based conservation measures. Gland, Switzerland. Available from https://portals.iucn.org/library/sites/library/files/documents/PATRS-003-En.pdf (accessed February 21, 2023).

Iwamura T, le Polain de Waroux Y, Mascia MB. 2018. Considering people in systematic conservation planning: insights from land system science. Frontiers in Ecology and the Environment 16:388–396.

Jonas HD, Bingham HC, Bennett NJ, Woodley S, Zlatanova R, Howland E, Belle E, Upton J, Gottlieb B, Kamath V, Lessmann J, Delli G, Dubois G, Ahmadia G, Claudet J, Cook C, Deza J, Grorud-Colvert K, Gurney G, Lemieux CJ and Ruiz L. 2024 Global status and emerging contribution of other effective area-based conservation measures (OECMs) towards the ‘30×30’ biodiversity Target 3. Front. Conserv. Sci. 5:1447434. doi: 10.3389/fcosc.2024.1447434

Joppa LN, Pfaff A. 2009. High and Far: Biases in the Location of Protected Areas. PLOS ONE 4:e8273. Public Library of Science.

Joseph LN, Maloney RF, Possingham HP. 2009. Optimal Allocation of Resources among Threatened Species: a Project Prioritization Protocol. Conservation Biology 23:328–338.

Kaim A, Watts ME, Possingham HP. 2017. On which targets should we compromise in conservation prioritization problems? Methods in Ecology and Evolution 8:1858–1865.

KBA Partnership. 2025. Who is using KBAs? Donors. Available from: https://www.keybiodiversityareas.org/who-using-kbas/donors (accessed November 14, 2025)

KBA Standards and Appeals Committee. 2022. Guidelines for using a global standard for the identification of Key Biodiversity Areas : version 1.1. IUCN, Gland, Switzerland. Available from https://portals.iucn.org/library/node/49131 (accessed February 12, 2023).

Kraus D, Enns A, Hebb A, Murphy S, Drake DAR, Bennett B. 2023. Prioritizing nationally endemic species for conservation. Conservation Science and Practice 5:e12845.

Liang J et al. 2016. Positive biodiversity-productivity relationship predominant in global forests. Science 354:aaf8957. American Association for the Advancement of Science.

Lim DYH, Starnes T, Plumptre AJ. 2023. Global priorities for biodiversity conservation in the United Kingdom. Biological Conservation 277:109798.

Lomba A, Pellissier L, Randin C, Vicente J, Moreira F, Honrado J, Guisan A. 2010. Overcoming the rare species modelling paradox: A novel hierarchical framework applied to an Iberian endemic plant. Biological Conservation 143:2647–2657.

Malatesta, L, Alves, T, Attorre, F, Brito, D, Cianciullo, S, Datizua, C, De Abreu, D, De Felici, S, De Sousa, C, Langa, C and Mate, B, 2023. BioNoMo: the biodiversity network of Mozambique. Rendiconti Lincei. Scienze Fisiche e Naturali, 34(1), pp.69–80.

Maron, M., Brownlie, S., Bull, J.W., Evans, M.C., von Hase, A., Quétier, F., Watson, J.E. and Gordon, A.. 2018. The many meanings of no net loss in environmental policy. Nature Sustainability, 1(1), pp.19–27.

Marchese C. 2015. Biodiversity hotspots: A shortcut for a more complicated concept. Global Ecology and Conservation 3:297–309.

Margules CR, Pressey RL. 2000. Systematic conservation planning. Nature 405:243–253. Nature Publishing Group.

Meyer C, Kreft H, Guralnick R, Jetz W. 2015. Global priorities for an effective information basis of biodiversity distributions. Nature Communications 6:8221. Nature Publishing Group.

MIMAIP, 2021. Elaboração do Plano de Ordenamento do Espaço Marítimo (POEM). Volume 1 - Proposta de Ordenamento do Espaço Marítimo, Tomo 5 – Processo de Participação Pública. Ministério do Mar, Águas Interiores e Pescas (MIMAIP), TPF, Biodesign. Rekatório final.

MITADER, 2015. National Strategy and Action Plan of Biological Diversity of Mozambique (2015-2035). Ministério da Terra, Ambiente e Desenvolvimento Rural (MITADER).

MITADER, 2019a. Sixth National report on the implementation of Convention on Biological Diversity in Mozambique. Page 152. Ministry of Land, Environment and Rural Development, Maputo. Available from https://www.cbd.int/doc/nr/nr-06/mz-nr-06-en.pdf (accessed February 28, 2023).

MITADER, 2019b. Plano nacional de desenvolvimento territorial. Relatório R.IV/02. Relatório de Avaliacao Ambiental e Social Estratégica. Versão Final.

Monadjem A, Schoeman MC, Reside A, Pio DV, Stoffberg S, Bayliss J, Cotterill FPD (Woody), Curran M, Kopp M, Taylor PJ. 2010. A recent inventory of the bats of Mozambique with documentation of seven new species for the country. Acta Chiropterologica 12:371–391.

Moore JL, Hauser CE, Bear JL, Williams NSG, McCarthy MA. 2011. Estimating detection— effort curves for plants using search experiments. Ecological Applications 21:601–607. Ecological Society of America.

Mori AS, Lertzman KP, Gustafsson L. 2017. Biodiversity and ecosystem services in forest ecosystems: a research agenda for applied forest ecology. Journal of Applied Ecology 54:12–27.

Neves IQ, Mathias M da L, Bastos-Silveira C. 2019. Mapping Knowledge Gaps of Mozambique’s Terrestrial Mammals. Scientific Reports 9:18184. Nature Publishing Group.

Nogueira JG et al. 2021. Alarming decline of freshwater trigger species in western Mediterranean key biodiversity areas. Conservation Biology 35:1367–1379.

Oppel S et al. 2018. Spatial scales of marine conservation management for breeding seabirds. Marine Policy 98:37–46.

Pereira M, Litulo C, Santos R, Costa Leal M, Fernandes R, Tibirica Y, Williams J, Atanassov B, Carreira F, Massingue A, Marques da Silva I. 2014. Mozambique marine ecosystems review. 10.13140/2.1.2092.5766.

Perino A et al. 2022. Biodiversity post-2020: Closing the gap between global targets and national-level implementation. Conservation Letters 15:12848.

Plumptre AJ, Ayebare S, Behangana M, et al. 2019. Conservation of vertebrates and plants in Uganda: Identifying Key Biodiversity Areas and other sites of national importance. Conservation Science and Practice. 2019;1:e7. 10.1002/csp2.7.

R Core Team. 2023. R: A language and environment for statistical computing. Vienna: R Foundation for Statistical Computing.

QGIS Development Team. 2023. QGIS Geographic Information System. Open Source Geospatial Foundation.

Rice WS, Sowman MR, Bavinck M. 2020.Using Theory of Change to improve post-2020 conservation: A proposed framework and recommendations for use. Conservation Science and Practice 2:e301.

Richards SL, Farooq H, Matimele H, Alves T, Datizua C, Langa C, Massingue A, Osborne J, Rokni S, de Sousa C, Darbyshire I. 2023. Identifying Mozambique’s most critical areas for plant conservation: An evaluation of protected areas and Important Plant Areas. Biotropica 55: 1183 – 1194. 10.1111/btp.13265

Rivers MC, Taylor L, Brummitt NA, Meagher TR, Roberts DL, Lughadha EN. 2011. How many herbarium specimens are needed to detect threatened species? Biological Conservation 144:2541–2547.

Roberts DL, Taylor L, Joppa LN. 2016. Threatened or Data Deficient: assessing the conservation status of poorly known species. Diversity and Distributions 22:558–565.

SANBI, 2024. Frequently Asked Questions About Key Biodiversity Areas in South Africa. Version: June 2024. Available from: https://www.sanbi.org/wp-content/uploads/2024/06/2024-06-SAKBA-FAQ.pdf (accessed January 12, 2023).

Schaaf T, Clamote Rodrigues D. 2016. Managing MIDAs: Harmonising the management of MultiInternationally Designated Areas: Ramsar sites, World Heritage sites, Biosphere Reserves and UNESCO Global Geoparks. IUCN, Gland, Switzerland. Available from https://portals.iucn.org/library/node/46176 (accessed February 23, 2023).

Smith RJ, Bennun L, Brooks TM, Butchart SH, Cuttelod A, Di Marco M, Ferrier, S, Fishpool LD, Joppa L, Juffe-Bignoli D and Knight AT. 2019. Synergies between the key biodiversity area and systematic conservation planning approaches. Conservation Letters, 12(1), p.e12625.

Stephenson PJ, Stengel C. 2020. An inventory of biodiversity data sources for conservation monitoring. PLOS ONE 15:e0242923.

Stephenson, PJ, Bowles-Newark, N, Regan, E, Stanwell-Smith, D, Diagana, M, Höft, R, Abarchi, H, Abrahamse, T, Akello, C, Allison, H and Banki, O. 2017. Unblocking the flow of biodiversity data for decision-making in Africa. Biological Conservation, 213, pp.335–340.

Thomas, CD and Gillingham, PK. 2015. The performance of protected areas for biodiversity under climate change. Biological Journal of the Linnean Society, 115(3), pp.718–730.

Tolley KA, Alexander GJ, Branch WR, Bowles P, Maritz B. 2016. Conservation status and threats for African reptiles. Biological Conservation 204:63–71.

Venter O et al. 2014. Targeting Global Protected Area Expansion for Imperiled Biodiversity. PLOS Biology 12:e1001891. Public Library of Science.

Venter O, Magrach A, Outram N, Klein CJ, Possingham HP, Di Marco M, Watson JEM. 2018. Bias in protected-area location and its effects on long-term aspirations of biodiversity conventions. Conservation Biology 32:127–134.

Visconti P, Butchart SHM, Brooks TM, Langhammer PF, Marnewick D, Vergara S, Yanosky A, Watson JEM. 2019. Protected area targets post-2020. Science 364:239–241. American Association for the Advancement of Science.

Wauchope HS et al. 2022. Protected areas have a mixed impact on waterbirds, but management helps. Nature 605:103–107. Nature Publishing Group.

WCS, Government of Mozambique, USAID. 2021a. Key Biodiversity Areas (KBAs) identified in Mozambique: Factsheets VOL. II. Red List of threatened species and ecosystems, identification and mapping of key biodiversity areas (KBAs) in Mozambique. Available from: https://comboprogram.org/Portals/1/Documents/1622457927-2021_KBAs_Moz_vol_ii_Factsheets_EN.pdf?ver=F2plHBM5lZ2x3-c8UYcMoA%3D%3D (accessed January 22, 2023).

WCS, Governo de Moçambique, USAID. 2021b. Red List of threatened species, ecosystems, identification and mapping of Key Biodiversity Areas (KBAs) in Mozambique - Final report (vol. 1). Available from https://comboprogram.org/Portals/1/Documents/1622463839-2021_KBAs_Moz_vol_i_Final_Report_EN.pdf?ver=YwbJaBNvIElPWYWfAowXEw%3D%3D (accessed February 20, 2023).

Zhang J, Nielsen SE, Grainger TN, Kohler M, Chipchar T, Farr DR. 2014. Sampling Plant Diversity and Rarity at Landscape Scales: Importance of Sampling Time in Species Detectability. PLoS ONE 9:e95334.

