## Supplemental Table 1 for "Identifying key biodiversity areas in Mozambique for conservation planning and policy"

**Supporting Information**

**Table S1**: List of trigger species with the KBA criteria and sub-criteria summarised for each species.

| Taxon Group | Species | IUCN Red List | KBA criteria triggered |
|---|---|---|---|
| Amphibian | Arthroleptis francei | VU | A1b(iv) |
| Amphibian | Hyperolius spinigularis | VU | A1b(iv) |
| Amphibian | Hyperolius swynnertoni | LC | B1(iv) |
| Amphibian | Nothophryne baylissi | EN | A1a(iv) |
| Amphibian | Nothophryne inagoensis | EN | A1a(iv); B1(iv) |
| Amphibian | Nothophryne ribauensis | EN | A1a(iv); A1e(iv); B1(iv) |
| Amphibian | Nothophryne unilurio | CR | A1a(iv); A1e(iv); B1(iv) |
| Amphibian | Strongylopus rhodesianus | VU | A1b(iv) |
| Bird | Apalis lynesi | NT | B1(i) |
| Bird | Arcanator orostruthus | VU | A1b(i) |
| Bird | Artisornis sousae | EN | A1a(i); A1e(i); B1(i) |
| Bird | Balearica regulorum | EN | A1a(i); A1c(i) |
| Bird | Bugeranus carunculatus | VU | A1b(i); A1d(i) |
| Bird | Chamaetylas choloensis | VU | A1b(i); B1(i); B2(i) |
| Bird | Gyps africanus | CR | A1a(i); A1c(i) |
| Bird | Hirundo atrocaerulea | VU | A1b(i) |
| Bird | Sternula balaenarum | VU | A1b(i); B1(i) |
| Bird | Swynnertonia swynnertoni | VU | A1b(i); B2(i) |
| Bird | Trigonoceps occipitalis | CR | A1a(i); A1c(i) |
| Fish (freshwater) | Chetia brevis | EN | A1a(iv) |
| Fish (freshwater) | Nothobranchius hengstleri | EN | A1a(iv); B1(iv) |
| Fish (freshwater) | Nothobranchius kadleci | NT | B1(iv) |
| Fish (freshwater) | Nothobranchius krammeri | VU | A1b(iv); B1(iv) |
| Fish (freshwater) | Nothobranchius krysanovi | VU | A1b(iv); B1(iv) |

| | | | |
|---|---|---|---|
| Fish (freshwater) | Nothobranchius niassa | VU | A1b(iv); B1(iv) |
| Fish (freshwater) | Oreochromis mortimeri | CR | A1a(iv); A1c(iv) |
| Fish (freshwater) | Silhouettea sibayi | EN | A1a(iv) |
| Fish (marine) | Mobula alfredi | VU | D1b(i) |
| Fish (marine) | Mobula birostris | EN | D1b(i) |
| Fish (marine) | Parablennius lodosus | VU | A1b(iv) |
| Fish (marine) | Rhincodon typus | EN | D1b(i) |
| Fish (marine) | Stegostoma tigrinum | EN | D1b(i) |
| Fish (marine) | Caranx Ignobilis | LC | D1b(i) |
| Insect | Acraea dondoensis | | B1(v) |
| Insect | Alaena lamborni | | B1(v) |
| Insect | Amblyphymus adspersus | NT | B1(iv) |
| Insect | Baliochila woodi | | B1(v) |
| Insect | Cymothoe baylissi | | B1(v) |
| Insect | Graphium junodi | | B1(v) |
| Insect | Iolaus (Epamera) malaikae | | B1(v) |
| Insect | Iolaus lulua | VU | A1b(v); B1(v) |
| Insect | Neocoenyra bioculata | | B1(v) |
| Insect | Pseudonympha cyclops | | B1(iv) |
| Insect | Scarabaeus bornemizzai | NT | B1(iv) |
| Mammal | Carpitalpa arendsi | VU | A1b(iv); B1(iv) |
| Mammal (marine) | Dugong dugon | VU | A1d(i) |
| Mammal | Hippopotamus amphibius | VU | A1b(i); A1d(i) |
| Mammal | Loxodonta africana | EN | A1a(i); A1c(i) |
| Mammal | Lycaon pictus | EN | A1a(i) |
| Mammal | Panthera leo | VU | A1b(i); A1d(i) |
| Mammal | Paraxerus vincenti | EN | A1a(iv) |
| Mammal (marine) | Sousa plumbea | EN | A1a(iv); A1c(iv) |
| Plant | Acacia latispina | VU | A1b(iv); B1(iv); B2(iv) |
| Plant | Acacia quiterajoensis | LC | B1(v) |
| Plant | Acacia torrei | LC | B1(iv) |

| | | | |
|---|---|---|---|
| Plant | Aeschynomene aphylla | VU | A1b(iv); B1(iv) |
| Plant | Agelanthus patelii | EN | A1a(v); A1e(v); B1(v) |
| Plant | Alloeochaete namuliensis | VU | A1b(iv); B1(iv) |
| Plant | Allophylus torrei | EN | A1a(v); B1(v) |
| Plant | Aloe plowesii | VU | A1b(v); B1(v) |
| Plant | Aloe ribauensis | EN | A1a(v); B1(v) |
| Plant | Aloe rulkensii | CR | A1a(i); A1e(i); B1(i) |
| Plant | Aspidoglossum glabellum | EN | A1a(v); B1(v) |
| Plant | Brachystelma nutans | VU | A1b(v); B1(v) |
| Plant | Buchnera subglabra | VU | A1b(v); B1(v) |
| Plant | Centella obtriangularis | VU | A1b(v); B1(v) |
| Plant | Chassalia colorata | EN | A1a(v); B1(v) |
| Plant | Cissus aristolochiifolia | VU | A1b(v); B1(v) |
| Plant | Clerodendrum lutambense | VU | A1b(iv); B1(iv) |
| Plant | Coffea schliebenii | VU | A1b(iv); B1(iv) |
| Plant | Coleus namuliensis | LC | B1(v) |
| Plant | Cordia stuhlmannii | VU | A1b(v); B1(v) |
| Plant | Crossopetalum mossambicense | EN | A1e(v) |
| Plant | Crotalaria insignis | VU | A1b(v); B1(v) |
| Plant | Crotalaria namuliensis | LC | B1(v) |
| Plant | Croton aceroides | EN | A1a(v); B1(v) |
| Plant | Croton Inhambanensis | VU | A1b(v); B1(v) |
| Plant | Cynanchum oresbium | VU | A1b(v); B1(v) |
| Plant | Danthoniopsis chimanimaniensis | EN | A1a(v); B1(v) |
| Plant | Didymosalpinx callianthus | EN | A1a(iv); B1(iv); B2(iv) |
| Plant | Dierama inyangense | EN | A1a(v); B1(v) |
| Plant | Disa zimbabweensis | VU | A1b(v); B1(v) |
| Plant | Dissotis pulchra | VU | A1b(v); B1(v) |
| Plant | Dissotis swynnertonii | VU | A1b(v); B1(v) |
| Plant | Dombeya leachii | EN | A1a(v); A1e(v); B1(v) |
| Plant | Dorstenia zambesiaca | VU | A1b(iv); B1(iv) |
| Plant | Ecbolium hastatum | EN | A1a(v); B1(v) |
| Plant | Empogona jenniferae | EN | A1a(v); B1(v) |
| Plant | Empogona maputensis | EN | A1a(v); B1(v) |

| | | | |
|---|---|---|---|
| Plant | Erica lanceolifera | VU | A1b(v); B1(v) |
| Plant | Faurea racemosa | EN | A1a(v); B1(v) |
| Plant | Ficus muelleriana | EN | A1a(v); B1(v) |
| Plant | Gladiolus zimbabweensis | VU | A1b(v); B1(v) |
| Plant | Grewia limae | EN | A1a(iv); B1(iv) |
| Plant | Gutenbergia westii | VU | A1b(v); B1(v) |
| Plant | Helixanthera schizocalyx | EN | A1a(v); B1(v) |
| Plant | Icuria dunensis | EN | A1a(v); B1(v) |
| Plant | Impatiens psychadelphoides | VU | A1b(v); B1(v) |
| Plant | Impatiens wuerstenii | VU | A1b(v); B1(v) |
| Plant | Isoglossa namuliensis | CR | A1a(v); A1e(v); B1(v) |
| Plant | Jatropha subaequiloba | VU | A1b(v); B1(v) |
| Plant | Justicia gorongozana | LC | B1(v) |
| Plant | Maerua andradae | LC | B1(iv) |
| Plant | Memecylon incisilobum | CR | A1a(i); A1e(i); B1(i) |
| Plant | Memecylon nubigenum | EN | A1a(v); B1(v) |
| Plant | Micklethwaitia carvalhoi | VU | A1b(iv); B2(iv) |
| Plant | Mimosa busseana | LC | B1(iv); B2(iv) |
| Plant | Morella chimanimaniana | EN | A1a(v); B1(v) |
| Plant | Neobolusia ciliata | EN | A1a(v); B1(v) |
| Plant | Ochna dolicharthros | VU | A1b(v); A1e(v); B1(v) |
| Plant | Olinia chimanimani | EN | A1a(v); B1(v) |
| Plant | Ormocarpum schliebenii | NT | B1(iv) |
| Plant | Otiophora lanceolata | VU | A1b(v); B1(v) |
| Plant | Oxyanthus biflorus | EN | A1a(iv); B1(iv); B2(iv) |
| Plant | Oxyanthus strigosus | EN | A1a(iv); A1e(iv); B1(iv); B2(iv) |
| Plant | Ozoroa gomesiana | VU | A1b(v); B1(v) |
| Plant | Pavetta gurueensis | VU | A1b(v); B1(v) |
| Plant | Pavetta lindina | EN | A1a(iv); B1(iv); B2(iv) |
| Plant | Plectranthus cucullatus | VU | A1b(v); B1(v) |
| Plant | Plectranthus mandalensis | VU | A1b(v); B1(v) |
| Plant | Polygala francisci | VU | A1b(v); B1(v) |
| Plant | Polygala zambesiaca | VU | A1b(v); B1(v) |
| Plant | Polysphaeria ribauensis | EN | A1a(v); A1e(v); B1(v) |
| Plant | Protea enervis | VU | A1b(v); B1(v) |

| | | | |
|---|---|---|---|
| Plant | Pseudomussaenda mozambicensis | EN | A1a(iv); B1(iv) |
| Plant | Psydrax fragrantissima | NT | B1(v) |
| Plant | Pterocephalus centennii | CR | A1a(v); A1e(v); B1(v) |
| Plant | Raphia australis | VU | A1b(i); B1(i) |
| Plant | Rhynchosia chimanimaniensis | EN | A1a(v); B1(v) |
| Plant | Rytigynia torrei | EN | A1a(v); B1(v) |
| Plant | Schistostephium oxylobum | VU | A1b(v); B1(v) |
| Plant | Schizochilus lepidus | VU | A1b(v); B1(v) |
| Plant | Sclerochiton apiculatus | VU | A1b(v); B1(v) |
| Plant | Sclerochiton hirsutus | VU | A1b(v); B1(v) |
| Plant | Sericanthe chimanimaniensis | VU | A1b(v); B1(v) |
| Plant | Streptocarpus brachynema | EN | A1a(v); A1e(v); B1(v) |
| Plant | Streptocarpus hirticapsa | VU | A1b(v); B1(v) |
| Plant | Streptocarpus myoporoides | EN | A1a(i); B1(i) |
| Plant | Streptocarpus umtaliensis | LC | B1(v) |
| Plant | Stylochaeton tortispathus | VU | A1b(iv); B1(iv) |
| Plant | Syncolostemon oritrephes | VU | A1b(v); B1(v) |
| Plant | Synsepalum sp. nov. | EN | A1a(v); B1(v) |
| Plant | Tarenna longipedicellata | VU | A1b(v); B1(v) |
| Plant | Tarenna pembensis | EN | A1a(iv)B1(iv) |
| Plant | Vepris allenii | EN | A1a(v); B1(v) |
| Plant | Vepris drummondii | VU | A1b(v); B1(v) |
| Plant | Vepris macedoi | EN | A1a(v); B1(v) |
| Plant | Warneckea cordiformis | CR | A1a(iv); A1e(iv); B1(iv) |
| Plant | Warneckea parvifolia | EN | A1a(v); B1(v) |
| Plant | Xerophyta kirkii | LC | B1(v) |
| Plant | Xerophyta pseudopinifolia | LC | B1(v) |
| Plant | Xylia mendoncae | VU | A1b(v); B1(v) |
| Plant | Xylopia torrei | EN | A1a(v); B1(v) |
| Plant | Xyris asterotricha | VU | A1b(v); B1(v) |
| Reptile | Afroedura gorongosa | LC | B1(iv) |
| Reptile | Atheris mabuensis | EN | A1a(iv); B1(iv); B2(iv) |
| Reptile | Cordylus meculae | LC | B1(iv) |
| Reptile | Cycloderma frenatum | EN | A1a(iv) |

| | | | |
|---|---|---|---|
| Reptile | Dipsadoboa montisilva | NT | B1(iv); B2(iv) |
| Reptile | Lygodactylus chiperone | | B1(iv); B2(iv) |
| Reptile | Lygodactylus inago | | B1(iv); B2(iv) |
| Reptile | Lygodactylus mabu | | B1(iv); B2(iv) |
| Reptile | Lygodactylus ribaue | | B1(iv); B2(iv) |
| Reptile | Mochlus lanceolatus | LC | B1(iv) |
| Reptile | Nadzikambia baylissi | NT | B1(iv); B2(iv) |
| Reptile | Nadzikambia chiperone | | B1(iv); B2(iv) |
| Reptile | Nadzikambia inago | | B1(iv); B2(iv) |
| Reptile | Nadzikambia namuli | | B1(iv); B2(iv) |
| Reptile | Nadzikambia ribaue | | B1(iv); B2(iv) |
| Reptile | Rhampholeon bruessoworum | CR | A1a(iv); B1(iv); B2(iv) |
| Reptile | Rhampholeon gorongosae | EN | A1a(iv); A1e(iv); B1(iv) |
| Reptile | Rhampholeon marshalli | VU | A1b(iv); B1(iv) |
| Reptile | Rhampholeon maspictus | NT | B1(iv); B2(iv) |
| Reptile | Rhampholeon namuli | | B1(iv); B2(iv) |
| Reptile | Rhampholeon nebulauctor | VU | A1b(iv); B1(iv); B2(iv) |
| Reptile | Rhampholeon tilburyi | EN | A1a(iv); B1(iv); B2(iv) |
| Reptile | Scelotes duttoni | LC | B1(iv) |
| Reptile | Scelotes insularis | LC | B1(iv) |
| Reptile | Scolecoseps broadleyi | NT | B1(iv) |
| Reptile | Smaug mossambicus | LC | B1(iv) |

**Table S2**: List of the 29 KBAs identified in Mozambique and the species that triggered KBA status per site. Of the 29 KBAs, 25 are terrestrial and four are marine namely: Vamizi, Great Bazaruto, Tofo and Ponta do Ouro Partial Marine Reserve.

| ID | KBA name | Province | KBA area km2 | KBA criteria | KBA trigger species | Taxonomic Group | IUCN status |
|---|---|---|---|---|---|---|---|
| 1 | Njesi Plateau | Niassa | 1996 | A1a(i); A1e(i); B1(i) | *Artisornis sousae* | Bird | EN |
| | | | | B1(iv) | *Cordylus meculae* | Reptile | LC |
| 2 | Niassa Special Reserve | Niassa | 42708 | A1a(i); A1c(i) | *Loxodonta africana* | Mammal | EN |
| | | | | A1b(i); A1d(i) | *Hippopotamus amphibius* | Mammal | VU |
| | | | | A1a(i) | *Lycaon pictus* | Mammal | EN |
| | | | | A1b(i); A1d(i) | *Panthera leo* | Mammal | VU |
| | | | | A1a(iv) | *Cycloderma frenatum* | Reptile | EN |
| | | | | B1(iv) | *Cordylus meculae* | Reptile | LC |
| | | | | A1b(iv); B1(iv) | *Nothobranchius niassa* | Freshwater Fish | VU |
| 3 | Palma | Cabo Delgado | 4556 | B1(iv) | *Scolecoseps broadleyi* | Reptile | NT |
| | | | | A1b(iv); B1(iv) | *Nothobranchius krammeri* | Freshwater Fish | VU |
| | | | | A1a(iv); B1(iv) | *Nothobranchius hengstleri* | Freshwater Fish | EN |
| | | | | A1a(v); B1(v) | *Chassalia colorata* | Plant | EN |
| | | | | A1a(iv); B1(iv) | *Grewia limae* | Plant | EN |
| | | | | A1a(iv); B1(iv); B2(iv) | *Pavetta lindina* | Plant | EN |

| | | | | | | | |
|---|---|---|---|---|---|---|---|
| | | | | A1e(v) | *Crossopetalum mossambicense* | Plant | EN |
| | | | | A1a(v); B1(v) | *Vepris allenii* | Plant | EN |
| | | | | A1b(iv); B1(iv) | *Clerodendrum lutambense* | Plant | VU |
| | | | | A1b(iv); B1(iv) | *Coffea schliebenii* | Plant | VU |
| | | | | A1a(iv); A1e(iv); B1(iv); B2(iv) | *Oxyanthus strigosus* | Plant | EN |
| | | | | B1(iv) | *Ormocarpum schliebenii* | Plant | NT |
| | | | | A1a(iv); B1(iv); B2(iv) | *Oxyanthus biflorus* | Plant | EN |
| | | | | A1b(v); A1e(v); B1(v) | *Ochna dolicharthros* | Plant | VU |
| | | | | B1(v) | *Acacia quiterajoensis* | Plant | LC |
| | | | | B1(iv); B2(iv) | *Mimosa busseana* | Plant | LC |
| | | | | A1a(iv); B1(iv); B2(iv) | *Didymosalpinx callianthus* | Plant | EN |
| 4 | Vamizi | Cabo Delgado | 87 | D1b(i) | *Caranx Ignobilis* | Marine Fish | LC |
| 5 | Quiterajo | Cabo Delgado | 3064 | A1a(iv); A1e(iv); B1(iv) | *Warneckea cordiformis* | Plant | CR |
| | | | | A1a(iv); B1(iv) | *Grewia limae* | Plant | EN |
| | | | | A1a(iv); A1e(iv); B1(iv); B2(iv) | *Oxyanthus strigosus* | Plant | EN |

| | | | | | | | |
|---|---|---|---|---|---|---|---|
| | | | | A1a(iv); B1(iv) | *Pseudomussaenda mozambicensis* | Plant | EN |
| | | | | A1a(iv); B1(iv); B2(iv) | *Pavetta lindina* | Plant | EN |
| | | | | A1a(v); B1(v) | *Vepris allenii* | Plant | EN |
| | | | | A1a(iv)B1(iv) | *Tarenna pembensis* | Plant | EN |
| | | | | A1b(iv); B2(iv) | *Micklethwaitia carvalhoi* | Plant | VU |
| | | | | A1b(iv); B1(iv) | *Stylochaeton tortispathus* | Plant | VU |
| | | | | A1b(iv); B1(iv); B2(iv) | *Acacia latispina* | Plant | VU |
| | | | | B1(v) | *Acacia quiterajoensis* | Plant | LC |
| | | | | B1(v) | *Justicia gorongozana* | Plant | LC |
| | | | | B1(iv) | *Maerua andradae* | Plant | LC |
| | | | | B1(iv); B2(iv) | *Mimosa busseana* | Plant | LC |
| | | | | B1(iv) | *Ormocarpum schliebenii* | Plant | NT |
| 6 | Taratibu | Cabo Delgado | 25 | A1a(iv); A1e(iv); B1(iv) | *Nothophryne unilurio* | Amphibian | CR |
| | | | | A1a(v); B1(v) | *Rytigynia torrei* | Plant | EN |
| 7 | Eráti | Nampula | 148 | A1a(v); B1(v) | *Allophylus torrei* | Plant | EN |
| 8 | Matibane Forest Reserve | Nampula | 109 | A1a(v); B1(v) | *Icuria dunensis* | Plant | EN |
| 9 | Ribáuè-Mphalwe | Nampula | 265 | B1(iv); B2(iv) | *Dipsadoboa montisilva* | Reptile | NT |

| | | | | | | | |
|---|---|---|---|---|---|---|---|
| | | | | A1a(iv); B1(iv); B2(iv) | *Rhampholeon tilburyi* | Reptile | EN |
| | | | | B1(iv); B2(iv) | *Nadzikambia ribaue* | Reptile | |
| | | | | B1(iv); B2(iv) | *Lygodactylus ribaue* | Reptile | |
| | | | | A1a(iv); A1e(iv); B1(iv) | *Nothophryne ribauensis* | Amphibian | EN |
| | | | | A1a(i); A1e(i); B1(i) | *Aloe rulkensii* | Plant | CR |
| | | | | A1a(v); B1(v) | *Aloe ribauensis* | Plant | EN |
| | | | | A1a(i); B1(i) | *Streptocarpus myoporoides* | Plant | EN |
| | | | | A1a(v); A1e(v); B1(v) | *Dombeya leachii* | Plant | EN |
| | | | | A1a(v); A1e(v); B1(v) | *Polysphaeria ribauensis* | Plant | EN |
| | | | | A1a(v); B1(v) | *Memecylon nubigenum* | Plant | EN |
| | | | | A1b(v); B1(v) | *Cynanchum oresbium* | Plant | VU |
| | | | | A1b(v); B1(v) | *Cissus aristolochiifolia* | Plant | VU |
| | | | | A1a(v); B1(v) | *Vepris macedoi* | Plant | EN |
| | | | | A1b(v); B1(v) | *Plectranthus cucullatus* | Plant | VU |
| 10 | Mount Inago | Nampula | 326 | A1a(iv); B1(iv); B2(iv) | *Rhampholeon bruessoworum* | Reptile | CR |
| | | | | B1(iv); B2(iv) | *Nadzikambia inago* | Reptile | |
| | | | | B1(iv); B2(iv) | *Lygodactylus inago* | Reptile | |
| | | | | A1a(iv); B1(iv) | *Nothophryne inagoensis* | Amphibian | EN |

| | | | | | | | |
|---|---|---|---|---|---|---|---|
| | | | | B1(v) | *Cymothoe baylissi* | Insect | |
| | | | | B1(v) | *Alaena lamborni* | Insect | |
| | | | | B1(v) | *Neocoenyra bioculata* | Insect | |
| 11 | APAIPS | Zambezia | 2512 | A1a(v); B1(v) | *Icuria dunensis* | Plant | EN |
| 12 | Mount Namuli | Zambezia | 53 | A1a(iv) | *Paraxerus vincenti* | Mammal | EN |
| | | | | B1(i) | *Apalis lynesi* | Bird | NT |
| | | | | A1b(i) | *Arcanator orostruthus* | Bird | VU |
| | | | | A1b(i); B1(i); B2(i) | *Chamaetylas choloensis* | Bird | VU |
| | | | | A1a(iv); B1(iv); B2(iv) | *Atheris mabuensis* | Reptile | EN |
| | | | | B1(iv); B2(iv) | *Nadzikambia namuli* | Reptile | |
| | | | | B1(iv); B2(iv) | *Rhampholeon namuli* | Reptile | |
| | | | | A1a(iv); B1(iv); B2(iv) | *Rhampholeon tilburyi* | Reptile | EN |
| | | | | A1b(iv) | *Arthroleptis francei* | Amphibian | VU |
| | | | | A1b(iv) | *Hyperolius spinigularis* | Amphibian | VU |
| | | | | A1a(iv) | *Nothophryne baylissi* | Amphibian | EN |
| | | | | B1(v) | *Baliochila woodi* | Insect | |
| | | | | B1(v) | *Cymothoe baylissi* | Insect | |
| | | | | B1(v) | *Iolaus (Epamera) malaikae* | Insect | |
| | | | | B1(v) | *Alaena lamborni* | Insect | |
| | | | | B1(v) | *Neocoenyra bioculata* | Insect | |

| | | | | | | | |
|---|---|---|---|---|---|---|---|
| | | | | A1a(v); A1e(v); B1(v) | *Agelanthus patelii* | Plant | EN |
| | | | | A1b(iv); B1(iv) | *Alloeochaete namuliensis* | Plant | VU |
| | | | | A1b(v); B1(v) | *Brachystelma nutans* | Plant | VU |
| | | | | B1(v) | *Coleus namuliensis* | Plant | LC |
| | | | | A1a(v); B1(v) | *Faurea racemosa* | Plant | EN |
| | | | | A1a(v); B1(v) | *Helixanthera schizocalyx* | Plant | EN |
| | | | | A1a(v); A1e(v); B1(v) | *Isoglossa namuliensis* | Plant | CR |
| | | | | A1a(v); B1(v) | *Memecylon nubigenum* | Plant | EN |
| | | | | A1b(v); B1(v) | *Pavetta gurueensis* | Plant | VU |
| | | | | A1b(v); B1(v) | *Plectranthus mandalensis* | Plant | VU |
| | | | | A1b(v); B1(v) | *Sclerochiton hirsutus* | Plant | VU |
| | | | | B1(v) | *Xerophyta kirkii* | Plant | LC |
| | | | | B1(v) | *Xerophyta pseudopinifolia* | Plant | LC |
| | | | | B1(v) | *Crotalaria namuliensis* | Plant | LC |
| 13 | Mount Mabu | Zambezia | 61 | A1b(i) | *Arcanator orostruthus* | Bird | VU |
| | | | | A1b(i); B1(i); B2(i) | *Chamaetylas choloensis* | Bird | VU |
| | | | | A1b(i); B2(i) | *Swynnertonia swynnertoni* | Bird | VU |
| | | | | A1a(iv); B1(iv); B2(iv) | *Atheris mabuensis* | Reptile | EN |
| | | | | B1(iv); B2(iv) | *Dipsadoboa montisilva* | Reptile | NT |
| | | | | B1(iv); B2(iv) | *Lygodactylus mabu* | Reptile | |

| | | | | | | | |
|---|---|---|---|---|---|---|---|
| | | | | B1(iv); B2(iv) | *Nadzikambia baylissi* | Reptile | NT |
| | | | | B1(iv); B2(iv) | *Rhampholeon maspictus* | Reptile | NT |
| | | | | A1b(iv) | *Arthroleptis francei* | Amphibian | VU |
| | | | | B1(v) | *Cymothoe baylissi* | Insect | |
| | | | | B1(v) | *Baliochila woodi* | Insect | |
| | | | | B1(v) | *Iolaus (Epamera) malaikae* | Insect | |
| | | | | B1(v) | *Neocoenyra bioculata* | Insect | |
| | | | | A1a(v); B1(v) | *Faurea racemosa* | Plant | EN |
| | | | | A1a(v); B1(v) | *Helixanthera schizocalyx* | Plant | EN |
| | | | | A1b(v); B1(v) | *Pavetta gurueensis* | Plant | VU |
| | | | | A1b(v); B1(v) | *Sclerochiton hirsutus* | Plant | VU |
| 14 | Mount Chiperone | Zambezia | 36 | A1b(i); B1(i); B2(i) | *Chamaetylas choloensis* | Bird | VU |
| | | | | A1b(iv); B1(iv); B2(iv) | *Rhampholeon nebulauctor* | Reptile | VU |
| | | | | B1(iv); B2(iv) | *Nadzikambia chiperone* | Reptile | |
| | | | | B1(iv); B2(iv) | *Lygodactylus chiperone* | Reptile | |
| 15 | Derre Forest Reserve | Zambezia | 3984 | A1b(iv); B1(iv) | *Nothobranchius krysanovi* | Freshwater Fish | VU |
| | | | | A1b(v); B1(v) | *Tarenna longipedicellata* | Plant | VU |
| 16 | Tchuma-Tchato | Tete | 38180 | A1b(i); A1d(i) | *Hippopotamus amphibius* | Mammal | VU |
| | Cahora Bassa Lake | | | A1a(iv); A1c(iv) | *Oreochromis mortimeri* | Freshwater Fish | CR |
| 17 | Serra Choa | Manica | 516 | A1b(i) | *Hirundo atrocaerulea* | Bird | VU |
| 18 | Machipanda | Manica | 756 | A1b(iv); B1(iv) | *Carpitalpa arendsi* | Mammal | VU |

| | | | | | | | |
|---|---|---|---|---|---|---|---|
| | | | | A1b(i) | *Hirundo atrocaerulea* | Bird | VU |
| 19 | Chimanimani National Park | Manica | 2371 | A1b(iv); B1(iv) | *Carpitalpa arendsi* | Mammal | VU |
| | | | | A1b(iv); B1(iv) | *Rhampholeon marshalli* | Reptile | VU |
| | | | | B1(iv) | *Smaug mossambicus* | Reptile | LC |
| | | | | B1(iv) | *Hyperolius swynnertoni* | Amphibian | LC |
| | | | | A1b(iv) | *Strongylopus rhodesianus* | Amphibian | VU |
| | | | | B1(iv) | *Pseudonympha cyclops* | Insect | |
| | | | | B1(v) | *Graphium junodi* | Insect | |
| | | | | A1a(v); A1e(v); B1(v) | *Pterocephalus centennii* | Plant | CR |
| | | | | A1a(v); B1(v) | *Danthoniopsis chimanimaniensis* | Plant | EN |
| | | | | A1a(v); B1(v) | *Morella chimanimaniana* | Plant | EN |
| | | | | A1a(v); B1(v) | *Neobolusia ciliata* | Plant | EN |
| | | | | A1a(v); B1(v) | *Olinia chimanimani* | Plant | EN |
| | | | | A1a(v); B1(v) | *Empogona jenniferae* | Plant | EN |
| | | | | A1a(v); B1(v) | *Ficus muelleriana* | Plant | EN |
| | | | | A1a(v); B1(v) | *Rhynchosia chimanimaniensis* | Plant | EN |
| | | | | A1a(v); B1(v) | *Aspidoglossum glabellum* | Plant | EN |
| | | | | A1a(v); B1(v) | *Dierama inyangense* | Plant | EN |
| | | | | A1a(v); B1(v) | *Synsepalum sp. nov.* | Plant | EN |

| | | | |
|---|---|---|---|
| A1b(iv); B1(iv) | *Aeschynomene aphylla* | Plant | VU |
| A1b(v); B1(v) | *Aloe plowesii* | Plant | VU |
| A1b(v); B1(v) | *Buchnera subglabra* | Plant | VU |
| A1b(v); B1(v) | *Centella obtriangularis* | Plant | VU |
| A1b(v); B1(v) | *Crotalaria insignis* | Plant | VU |
| A1b(v); B1(v) | *Dissotis pulchra* | Plant | VU |
| A1b(v); B1(v) | *Dissotis swynnertonii* | Plant | VU |
| A1b(v); B1(v) | *Erica lanceolifera* | Plant | VU |
| A1b(v); B1(v) | *Gutenbergia westii* | Plant | VU |
| A1b(v); B1(v) | *Otiophora lanceolata* | Plant | VU |
| A1b(v); B1(v) | *Protea enervis* | Plant | VU |
| A1b(v); B1(v) | *Schistostephium oxylobum* | Plant | VU |
| A1b(v); B1(v) | *Schizochilus lepidus* | Plant | VU |
| A1b(v); B1(v) | *Sericanthe chimanimaniensis* | Plant | VU |
| A1b(v); B1(v) | *Streptocarpus hirticapsa* | Plant | VU |
| A1b(v); B1(v) | *Syncolostemon oritrephes* | Plant | VU |
| A1b(v); B1(v) | *Vepris drummondii* | Plant | VU |
| A1b(v); B1(v) | *Xyris asterotricha* | Plant | VU |
| B1(v) | *Streptocarpus umtaliensis* | Plant | LC |
| A1b(v); B1(v) | *Polygala zambesiaca* | Plant | VU |
| A1b(v); B1(v) | *Impatiens psychadelphoides* | Plant | VU |
| A1b(v); B1(v) | *Gladiolus zimbabweensis* | Plant | VU |
| A1b(v); B1(v) | *Disa zimbabweensis* | Plant | VU |

| | | | | | | | |
|---|---|---|---|---|---|---|---|
| 20 | Gorongosa and Marromeu Complex | Sofala | 23088 | A1b(i); A1d(i) | *Hippopotamus amphibius* | Mammal | VU |
| | | | | A1a(i); A1c(i) | *Gyps africanus* | Bird | CR |
| | | | | A1a(i) | *Lycaon pictus* | Mammal | EN |
| | | | | A1a(i); A1c(i) | *Loxodonta africana* | Mammal | EN |
| | | | | A1b(i); A1d(i) | *Panthera leo* | Mammal | VU |
| | | | | A1a(i); A1c(i) | *Trigonoceps occipitalis* | Bird | CR |
| | | | | A1a(i); A1c(i) | *Balearica regulorum* | Bird | EN |
| | | | | A1b(i); A1d(i) | *Bugeranus carunculatus* | Bird | VU |
| | | | | B1(iv) | *Afroedura gorongosa* | Reptile | LC |
| | | | | A1a(iv) | *Cycloderma frenatum* | Reptile | EN |
| | | | | A1a(iv); A1e(iv); B1(iv) | *Rhampholeon gorongosae* | Reptile | EN |
| | | | | B1(iv) | *Nothobranchius kadleci* | Freshwater Fish | NT |
| | | | | B1(v) | *Acraea dondoensis* | Insect | |
| | | | | B1(v) | *Graphium junodi* | Insect | |
| | | | | B1(iv) | *Acacia torrei* | Plant | LC |
| | | | | A1b(v); B1(v) | *Impatiens wuerstenii* | Plant | VU |
| | | | | B1(v) | *Justicia gorongozana* | Plant | LC |
| | | | | A1a(v); A1e(v); B1(v) | *Streptocarpus brachynema* | Plant | EN |
| | | | | A1b(v); B1(v) | *Cordia stuhlmannii* | Plant | VU |
| | | | | A1b(iv); B1(iv) | *Dorstenia zambesiaca* | Plant | VU |
| | | | | A1b(v); B1(v) | *Tarenna longipedicellata* | Plant | VU |

| | | | | | | | |
|---|---|---|---|---|---|---|---|
| 21 | Inhassoro-Vilankulos | Inhambane | 5357 | A1b(v); B1(v) | *Croton Inhambanensis* | Plant | VU |
| | | | | A1a(v); B1(v) | *Ecbolium hastatum* | Plant | EN |
| | | | | A1a(v); B1(v) | *Croton aceroides* | Plant | EN |
| | | | | A1b(v); B1(v) | *Ozoroa gomesiana* | Plant | VU |
| | | | | A1b(v); B1(v) | *Xylia mendoncae* | Plant | VU |
| 22 | Great Bazaruto | Inhambane | 5236 | A1d(i) | *Dugong dugon* | Mammal | VU |
| | | | | A1a(iv); A1c(iv) | *Sousa plumbea* | Mammal | EN |
| | | | | B1(iv) | *Mochlus lanceolatus* | Reptile | LC |
| | | | | B1(iv) | *Scelotes insularis* | Reptile | LC |
| | | | | B1(iv) | *Scelotes duttoni* | Reptile | LC |
| | | | | A1b(v); B1(v) | *Jatropha subaequiloba* | Plant | VU |
| 23 | Tofo | Inhambane | 342 | D1b(i) | *Mobula alfredi* | Fish | VU |
| | | | | D1b(i) | *Mobula birostris* | Fish | EN |
| | | | | D1b(i) | *Rhincodon typus* | Fish | EN |
| | | | | D1b(i) | *Stegostoma tigrinum* | Fish | EN |
| 24 | Chongoene | Gaza | 33 | A1b(v); B1(v) | *Iolaus lulua* | Insect | VU |
| 25 | Manhiça-Bilene | Gaza | 2070 | A1a(iv) | *Chetia brevis* | Freshwater Fish | EN |
| | | | | A1a(i); A1e(i); B1(i) | *Memecylon incisilobum* | Plant | CR |
| | | | | A1b(i); B1(i) | *Raphia australis* | Plant | VU |
| 26 | Matutuine | Maputo | 195 | A1b(v); B1(v) | *Sclerochiton apiculatus* | Plant | VU |
| | | | | A1b(v); B1(v) | *Polygala francisci* | Plant | VU |

| | | | | | | | |
|---|---|---|---|---|---|---|---|
| 27 | Licuáti Forest Reserve | Maputo | 141 | A1b(v); B1(v) | *Sclerochiton apiculatus* | Plant | VU |
| | | | | A1b(v); B1(v) | *Polygala francisci* | Plant | VU |
| | | | | A1a(v); B1(v) | *Empogona maputensis* | Plant | EN |
| | | | | B1(v) | *Psydrax fragrantissima* | Plant | NT |
| | | | | A1a(v); B1(v) | *Warneckea parvifolia* | Plant | EN |
| | | | | A1a(v); B1(v) | *Xylopia torrei* | Plant | EN |
| 28 | Maputo National Park | Maputo | 1040 | A1b(v); B1(v) | *Sclerochiton apiculatus* | Plant | VU |
| | | | | B1(iv) | *Scarabaeus bornemizzai* | Insect | NT |
| | | | | A1a(iv) | *Silhouettea sibayi* | Freshwater Fish | EN |
| | | | | A1a(i); A1c(i) | *Loxodonta africana* | Mammal | EN |
| | | | | B1(iv) | *Amblyphymus adspersus* | Insect | NT |
| 29 | Ponta do Ouro Partial Marine Reserve | Maputo | 698 | D1b(i) | *Caranx Ignobilis* | Marine Fish | LC |
| | | | | A1b(iv) | *Parablennius lodosus* | Fish | VU |
| | | | | A1a(iv); A1c(iv) | *Sousa plumbea* | Mammal | EN |
